# Metabolic fate of human immunoactive sterols in *Mycobacterium tuberculosis*

**DOI:** 10.1101/2020.07.07.192294

**Authors:** Tatsiana Varaksa, Sergey Bukhdruker, Irina Grabovec, Egor Marin, Anton Kavaleuski, Anastasiia Gusach, Kirill Kovalev, Ivan Maslov, Aleksandra Luginina, Dmitry Zabelskiy, Roman Astashkin, Mikhail Shevtsov, Sviatlana Smolskaya, Anna Kavaleuskaya, Polina Shabunya, Alexander Baranovsky, Vladimir Dolgopalets, Yury Charnou, Aleh Savachka, Raisa Litvinovskaya, Alakseij Hurski, Evgeny Shevchenko, Andrey Rogachev, Alexey Mishin, Valentin Gordeliy, Andrei Gabrielian, Darrell E. Hurt, Boris Nikonenko, Konstantin Majorov, Alexander Apt, Alex Rosenthal, Andrei Gilep, Valentin Borshchevskiy, Natallia Strushkevich

**Author notes:** these authors contributed equally to this work (Tatsiana Varaksa, Sergey Bukhdruker).

## Abstract

*Mycobacterium tuberculosis* (Mtb) infection is among top ten causes of death worldwide, and the number of drug-resistant strains is increasing. The direct interception of human immune signaling molecules by Mtb remains elusive, limiting drug discovery. Oxysterols and secosteroids regulate both innate and adaptive immune responses. Here we report a functional, structural, and bioinformatics study of Mtb enzymes initiating cholesterol catabolism and demonstrated their interrelation with human immunity. We show that these enzymes metabolize human immune oxysterol messengers. Rv2266 – the most potent among them – can also metabolize vitamin D3 (VD3) derivatives. High-resolution structures show common patterns of sterols binding and reveal a site for oxidative attack during catalysis. Finally, we designed a compound that binds and inhibits three studied proteins. The compound shows activity against Mtb H37Rv residing in macrophages. Our findings contribute to molecular understanding of suppression of immunity and suggest that Mtb has its own transformation system resembling the human phase I drug-metabolizing system.

## Introduction

*Mycobacterium tuberculosis* (Mtb) is one of the most ancient of known human pathogens. It has accompanied hominids for 70 thousand years. Consequently, Mtb has developed innate resistance to the human immune response (*1, 2*). Continuing spread of drug resistance to widely used therapeutics over recent decades has become a substantial clinical problem. In this regard, the identification of novel molecular drug targets is of pivotal importance.

Mtb has a powerful sterol-catabolizing system that is essential for its virulence (*3*). The enzymes involved in catabolism of the cholesterol of the host are receiving growing attention as promising new targets for anti-tuberculosis (anti-TB) drug discovery (*4, 5*). The 3β-hydroxysteroid dehydrogenase (3βHSD) Rv1106c and monooxygenases Rv3518c (CYP142), Rv3545c (CYP125), and Rv2266 (CYP124) initiate cholesterol degradation in Mtb (*6, 7*). *In vitro* experiments with knockout mutants demonstrated that 3βHSD is required for the growth of Mtb on cholesterol as the sole carbon source (*6*). The inability of a 3βHSD transposon mutant to grow on cholesterol indicates that 3βHSD is the sole enzyme oxidizing the 3β-hydroxyl moiety of cholesterol in Mtb. CYP125 catalyzes the terminal hydroxylation of cholesterol and is required for *in vivo* survival of Mtb (*8*). Knockout of this gene significantly reduces the development of Mtb *in vivo* at the very first stages of infection (*8*). CYP142 serves as a functionally redundant analog for CYP125 (*9*). CYP124 can also metabolize cholesterol (*7*). However, it is not essential for *in vitro* growth on cholesterol (*10*), indicating that cholesterol catabolism is not a main function of this enzyme. CYP124 was suggested to perform hydroxylation of methyl-branched lipids (*11*), VD3, and 7-dehydrocholesterol (7DHC) (*12*) based on *in vitro* assays with the purified enzyme. Considering that cholesterol is not an essential source of nutrition for Mtb during infection, the physiological function of these enzymes could extend beyond cholesterol catabolism.

Cholesterol derivatives, such as oxysterols and VD3, are generally accepted as important factors of the human innate immune system – they directly regulate the inflammatory programming of macrophages (*13, 14*) and participate in the development of immune response (Table S1 summarizes known examples). We hypothesized that cholesterol metabolism by Mtb is not only a source of carbon and energy, but that it offers the pathogen a unique evolutionary ability to degrade immunoactive cholesterol derivatives. This ability of Mtb has not been evaluated before. Here we report that Mtb enzymes 3βHSD, CYP124, CYP125, and CYP142 can bind and metabolize a panel of human immunoactive oxysterols *in vitro*. We determined the binding constant and turnover number and identified the most important metabolic products. The most potent enzyme, CYP124, is active against both oxysterol and VD3 immunoactive derivatives. The putative relationship of the cholesterol-metabolizing enzymes of Mtb with human immune pathways found in our study is summarized in Fig. 1. Our further bioinformatic analysis confirms the importance of studied enzymes. Phylogenetic analysis shows that CYP124, CYP125, and 3βHSD are evolutionarily conserved in pathogenic mycobacteria, while transcriptome analysis suggests that CYP124 and CYP125 along with the mammalian-cell-entry proteins (mce) group together for the uptake and degradation of steroid molecules at early stages of infection. Considering the revealed importance of CYP124, we further solved its high-resolution structures in complex with substrates and inhibitors. Finally, we designed an inhibitor with an alternative fold that mimics sterol substrates and showed its effectiveness against Mtb residing in macrophages. All together our findings provide a structural basis for sterols recognition and modification by Mtb and advance current knowledge of host-pathogen interactions.

**Fig. 1.**
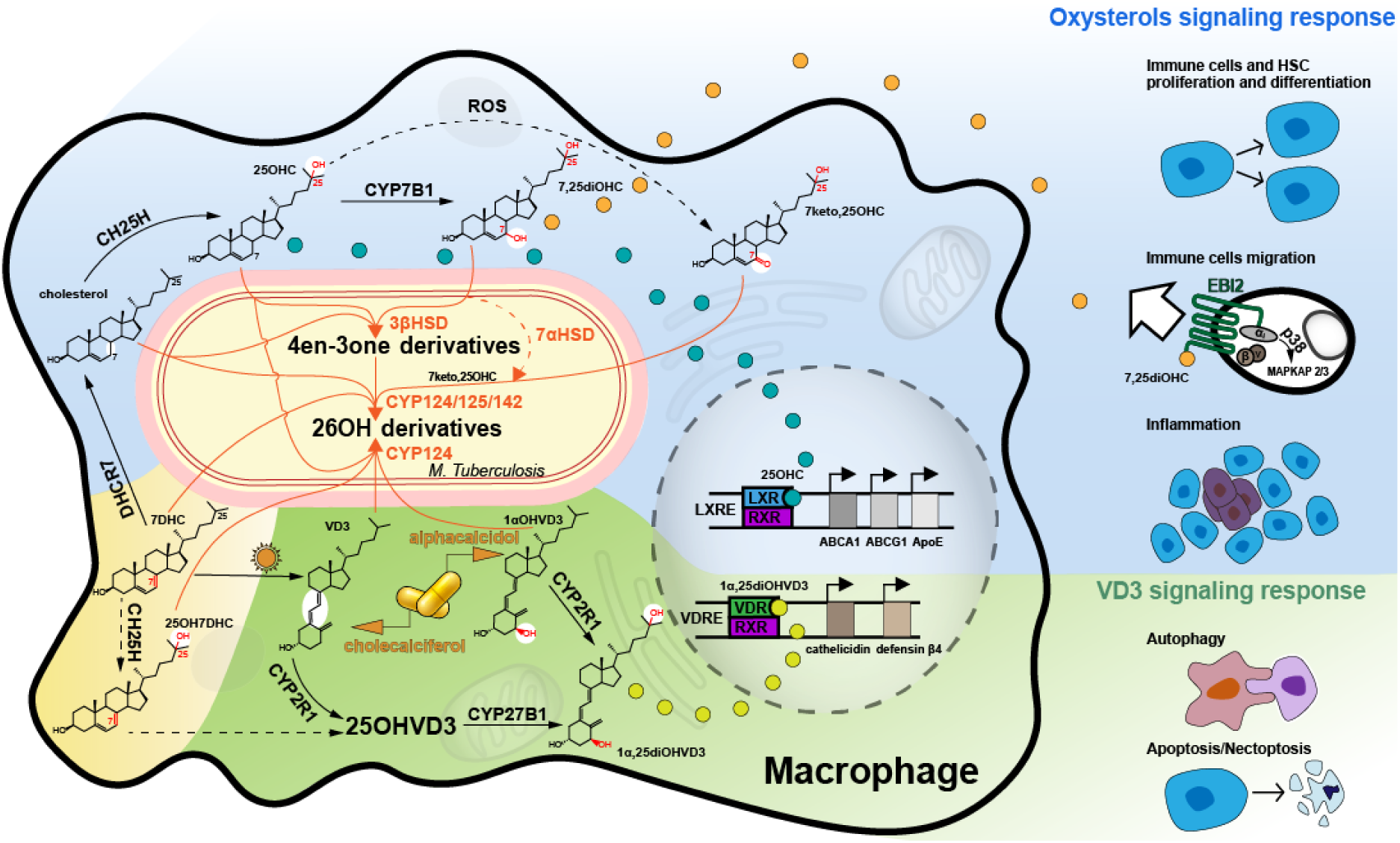
Oxysterol/secosterol pathways in human macrophages. A Mtb bacillus is shown inside a macrophage cell. Oxysterol transformation and signaling is shown inside and outside the macrophage on the blue background (see Discussion), VD3 activation and signaling on the green background, and 7DHC derivatives on the sand-colored background.

## Results

### CYP124, CYP125, CYP142 and 3βHSD metabolize immunoactive oxysterols *in vitro*

To test our idea that cholesterol-catabolizing enzymes process immune system messengers, we tested known immunoactive sterols, including side chain and B-ring derivatives, for being substrates for four cholesterol-metabolizing Mtb enzymes (3βHSD, CYP124, CYP125 and CYP142).

For the three CYPs, we performed a spectrophotometric titration assay to estimate the affinity of sterols to the active sites. Binding of sterols induce type I spectral response corresponding to substrate binding (Fig. S1, Fig. S2, and Fig. S3 for CYP124, CYP125, and CYP142, respectively). The calculated Kd values are within the micromolar range (Table S2). Notably, CYP124 is able to bind the majority of the tested compounds.

All of the selected sterols were further used as substrates for purified recombinant Mtb CYPs in activity assays (Table 1). Oxysterols with a hydroxyl group at the C25 atom, which did not induce typical spectral changes in CYPs (Table S2), were also tested in activity assays. 3βHSD was tested against oxysterols but not VD3 derivatives since no 3β-hydroxylation is not possible in secosteroids (Table 2).

**Table 1.**
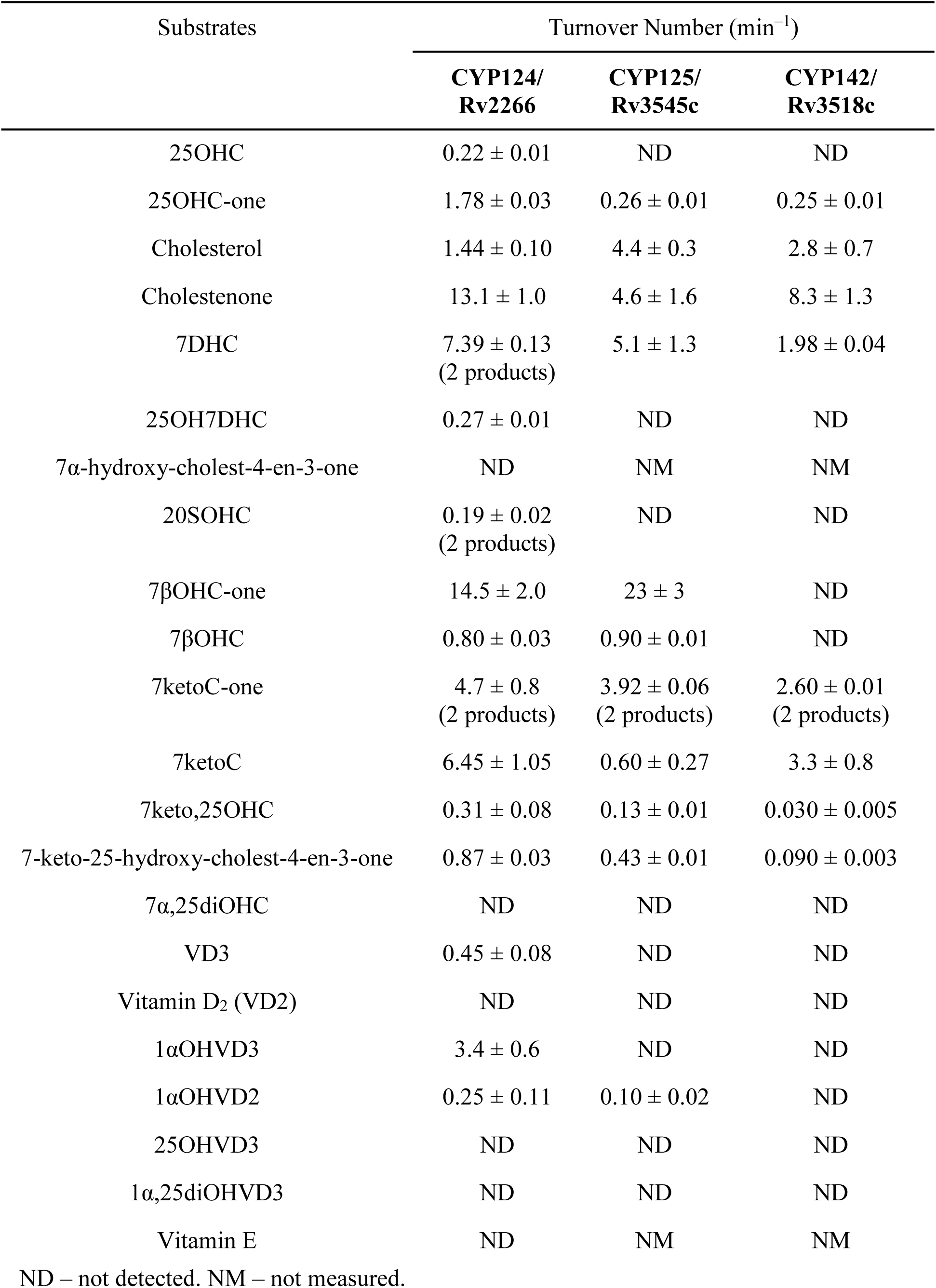
Catalytic activity of purified recombinant Mtb CYP124, CYP125, and CYP142.

**Table 2.** Catalytic activity of purified recombinant Mtb 3βHSD.

| Substrate | Turnover Number (min<sup>-1</sup>) |
| --- | --- |
| Cholesterol | 0.129 ± 0.004 |
| 25OHC | 0.218 ± 0.002 |
| 7βOHC | 0.688 ± 0.007 |
| 25OH7DHC | 1.31 ± 0.08 |
| 7ketoC | 0.06 ± 0.03 |
| 7keto,25OHC | ND |
| 7α,25diOHC | 1.09 ± 0.10 (major product),<br>0.165 ± 0.019 (minor product) |
| 17α-hydroxy-pregnenolone | 0.169 ± 0.002 |
| DHEA | 0.36 ± 0.07 |
| 5-androsten-3β,17β-diol | 0.053 ± 0.004 |

Among all the selected sterols, 25-hydroxy-cholesterol (25OHC) (Fig. 1) is of pivotal interest since it is a potent bioactive lipid in both the innate and adaptive immune systems (*15*). 25OHC is produced and secreted by macrophages in response to activation of Toll-like receptors (TLRs) or interferon receptors (INF) in early stages of infection (see Table S1 for more details). We found that 3βHSD catalyzes the conversion of 25OHC (Table 2) with formation of 25-hydroxy-cholest-4-en-3-one (25OHC-one), as confirmed by HPLC analysis. The product is further metabolized by all the studied CYPs, albeit with significantly different efficiencies (Table 1). CYP124 is the most active towards 25OHC-one and produces 25,26-dihydroxy-cholest-4-en-3-one (25,26diOHC-one), as determined by NMR spectroscopy (Fig. S4). Of note, CYP124 is the only one of the CYPs that direct metabolizes 25OHC. To confirm the possibility of the degradation of this oxysterol by mycobacteria, we assessed 25OHC metabolism by living Mtb cells. Whole-cell *in vitro* assay (see the Methods section) confirms 25OHC degradation by the Mtb H37Rv strain. Metabolic activity of Mtb cells estimated by substrate consumption is (1.03 ± 0.15) × 10^6^ molecules of 25OHC/Mtb cell/h.

Among all the tested oxysterols, 3βHSD has the highest activity toward 25-hydroxy-7-dehydrocholesterol (25OH7DHC), with a turnover number of 1.3 min^−1^ (Table 2). CYP124 metabolizes this endogenous human steroid as well (Table 3). Even though the role of 25OH7DHC in immunity is unexplored, similarly to 7DHC it can be a precursor of diverse oxysterols (*16*).

We also demonstrated that 3βHSD metabolizes 7α,25-dihydroxy-cholesterol (7α,25diOHC) (Fig. 1, Table 2), a natural agonist of the EBI2 receptor that is implicated in immune cell migration in humans (*17, 18*). In this reaction, 3βHSD acts as a ligand-degenerating enzyme (*19*), converting 7α,25diOHC to the inactive 3-one compound. Metabolism of 7α,25diOHC by the studied CYPs was not detected.

3βHSD is also active toward pro-inflammatory 7β-hydroxy-cholesterol (7βOHC) and produces 7β-hydroxy-cholest-4-en-3-one (7βOHC-one), which is further effectively metabolized by CYP124 and CYP125 to 7β,26-dihydroxy-cholest-4-en-3-one (Table 2, Table 1). 7βOHC can also be metabolized directly by CYP124 and CYP125, albeit at an extremely low rate. Interestingly, CYP142 is not active toward 7β-hydroxy-derivatives, but it efficiently metabolizes respective 7-keto-derivatives. Both 7-keto-cholesterol (7ketoC) and 7ketoC-one are also metabolized by CYP124. Moreover, the studied CYPs can hydroxylate 7keto,25OHC and 7-keto-25-hydroxy-cholest-4-en-3-one. The activity of all the studied enzymes with this group of oxysterols increases their pathophysiological relevance.

CYP124 is the only CYP that is able to bind and metabolize 20S-hydroxy-cholesterol (20SOHC, Table 1), a ligand of smoothened receptor (SMO) in the Hedgehog signaling pathway, which is also activated in response to pathogens, including mycobacteria (*20, 21*).

Thus, we found that the studied Mtb enzymes have a remarkably broad ability to metabolize bioactive sterols that are known as agonists of essential human receptors: a) 25OHC (agonist of Liver X receptor (LXR)); b) 7βOHC (agonist of orphan RORα/γ); c) 7α,25diOHC (agonist of the Epstein-Barr virus-induced G-protein coupled receptor 2 (EBI2)); d) 20SOHC (a ligand of SMO in the Hedgehog signaling pathway). The 3βHSD has a notable role since its activity increases the catalytic efficiency of CYPs with all the tested cholesterol derivatives.

### Structural basis for oxysterol hydroxylation

The above-mentioned findings indicate that the Mtb enzymes have broad spectra of abilities to degrade signaling oxysterol molecules of the host. Among the three selected CYPs, CYP124 showed the broadest and highest catalytic activity toward the tested oxysterol derivatives.

The binding of steroids to CYP125 and CYP142 was previously structurally exemplified in greater detail in the case of the cholestenone complex (PDB ID 2X5W and 2YOO) − at 1.58 Å and 1.69 Å resolution, correspondingly (*22, 23*), but no structure for the CYP124–sterol complex has been published so far. Here we complement the published data with the structure of the CYP124– cholestenone complex of the same quality of 1.65 Å resolution (Table S3).

The overall structure of the complex resembles the previously published structure of CYP124 in complex with phytanic acid (PDB ID 2WM4). The electron density for the ligand is unambiguously defined (Fig. S5A), showing the shape of a binding tunnel with 24 lining residues (depicted in Fig. 2A). The tunnel narrows towards the catalytic site to tightly enclose the aliphatic side-chain between BC-loop, αF and αI. It opens in the opposite direction to create more space for the tetracycle (see Fig. 2, F and K, for two cross-sections of the binding cavity).

**Fig. 2.**
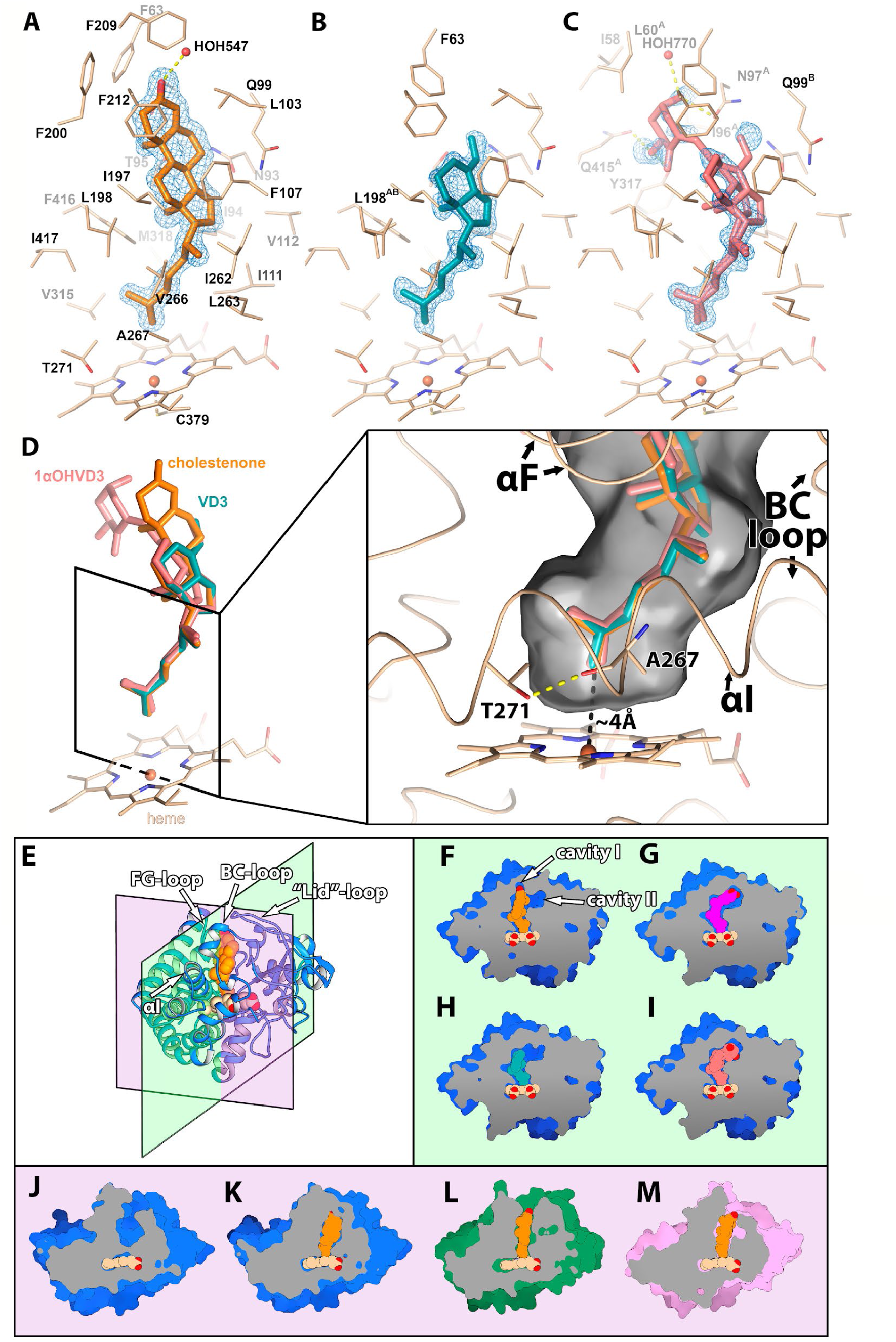
Structures of CYP124 in complexes with substrates. Structure of CYP124 binding pocket with substrates: cholestenone, **A;** VD3, **B;** and 1αOHVD3, **C**. Only side chains within 5 Å and directly H-bonded water molecules are shown for clarity. The *2mF*_*o*_−*DF*_*c*_ composite omit maps are contoured at 1σ. **D**, Superposition of CYP124 substrates in the active site. **E**, CYP124 secondary structure with two cross-sections presenting right (green) and bottom (lilac) panels. The green panel demonstrates CYP124 ligand-binding cavities for the following substrates: cholestenone, **F;** phytanic acid (PDB ID 2WM4), **G**; VD3, **H;** and 1αOHVD3, **I**. The lilac panel highlights ligand-binding cavities of apo CYP124 (PDB ID 2WM5), **J;** and the following complexes: CYP124–cholestenone, **K;** CYP125–cholestenone (PDB ID 2X5W), **L;** and CYP142– cholestenone (PDB ID 2YOO), **M**.

Compared to the apo-form (PDB ID 2WM5), the binding of cholestenone causes conformational changes in the proximity of the heme that are consistent with those of the phytanic acid complex (*11*). Water, coordinating the heme iron, is displaced from the active site, while the carbonyl of A267 rotates to form a hydrogen bond with the side-chain of T271, this resulting in the straightening of αI. Simultaneously, paired movement of the BC-loop with the FG-loop, as well as ordering of the “lid”-loop (Fig. 2E), leads to the enclosure of the active site from the solvent on the distal side.

There are notable differences in the binding modes of cholestenone in CYP124, CYP125 and CYP142 (see Fig. 2J-M). First, the steroid plane in the CYP124–cholestenone complex is rotated perpendicular to those in the two other proteins. Second, differences in positioning of the terminus of the isoprenoid chain above the heme are consistent with previously identified stereoisomeric products: (25R)-26-hydroxy configuration for CYP124 and CYP142 and (25S)-26-hydroxy configuration for CYP125 (*7*). Finally, the binding cavity remains opened in CYP125 and CYP142, exposing cholestenone A-ring to the solvent. In CYP124 structure, cholestenone is fully enclosed in the cavity. This implies that CYP125 and CYP142 may tolerate binding to more complex cholesterol A-ring derivatives, such as esters.

### Structural basis for hydroxylation of VD3 derivatives

Besides the actions of oxysterols, the innate and adaptive immune responses are modulated by VD3 and its derivatives (*24*). Previously, we detected the transformation of VD3 by a recombinant CYP124 (*12*); however, the identity of the product was not confirmed, and full spectra of potentially important VD3 derivatives were not evaluated in comparison with other cholesterol-metabolizing Mtb CYPs. Here, we further demonstrate that out of the three studied CYPs, only CYP124 binds VD3 derivatives. CYP124 also efficiently converts 1α-hydroxy-derivatives of both ergocalciferol (vitamin D2) and cholecalciferol (VD3) − 1α-hydroxy-vitamin D2 (1αOHVD2) and 1α-hydroxy-vitamin D3 (1αOHVD3, alfacalcidol), correspondingly, to their hydroxy products (Table 1). To get structural insight into the binding mode of host secosteroids, we solved structures of CYP124 in complex with VD3 and 1αOHVD3 with 1.3 Å and 1.2 Å resolution, respectively (Table S3). In both cases, the electron density unambiguously shows ligands in the active site of the protein (Fig. S5, B and C) with the C26 atom positioned for hydroxylation (Fig. 2, B and C). This type of modification is similar to the products profile of the mammalian C26 hydroxylation pathway of VD3 catabolism (*25*). The secosteroid A-ring is not resolved in the VD3 structure, probably due to its high conformational flexibility. In the CYP124−1αOHVD3 complex, the secosterol moiety of the ligand adopts two alternative conformations. One of them is virtually identical to the VD3 structure, while in the second an additional OH-group of the A-ring H-bonds with Q415 and Y317. In this case, the position of the A-ring is stabilized in cavity II, and the corresponding electron density is clearly visible. Surprisingly, addition of A-ring H-bonds does not improve the apparent dissociation constant Kd of VD3 compared with 1αOHVD3 (see Table S2); however, it may give a hint of a much more effective catabolism of 1αOHVD3 by CYP124 (Table 1).

Cholestenone and secosteroid substrates of CYP124 have almost identically positioned terminal side-chains surrounded by the residues from helixes F and I and the BC-loop region in the active site of the protein (fig. 2D). This indicates that methyl-branching at the ω and ω-4 positions is a crucial requirement for CYP124 substrates. But the surrounding of the secosteroid part is significantly different from that of cholestenone (Fig. 2, H and I). In the case of cholestenone, the binding pocket forms an extension − cavity I − to fully accommodate the rigid tetracycle moiety, and at the same time the sideward cavity II is filled with water molecules. In the case of more flexible secosteroids, cavity I is collapsed by a 4-Å movement of F63 and the A-ring extends into cavity II. The shape of the binding pocket in this case is very similar to that of the CYP124−phytanic acid complex (Fig. 2G). These observations provide a structural basis for the functional diversity of CYP124.

### Phylogenetic and bioinformatic transcriptome analysis of sterol-metabolizing Mtb enzymes

Phylogenetic reconstruction revealed that all three of the steroid-metabolizing CYPs are close relatives (Fig. 3 and Fig. S6 for more details). The CYP142 enzyme is in the first clade of steroid-metabolizing CYPs that diverged from the common ancestor. It is widespread in Mycobacteria, but it is not conservative in the *Mycobacterium tuberculosis* complex (MTC) − a group of Mycobacterium species that can cause tuberculosis (TB) in humans or other animals. We therefore presume that CYP142 is not significant for the development of TB. The CYP125 clade diverges after CYP142, whereas CYP124 is the most evolutionarily recent member of the steroid-catabolizing CYPs. CYP124, and CYP125 are both conserved in Mycobacteria, including the MTC group, which points to their importance for the development of Mtb. They have notably different species distribution though. The number of CYP125 paralogs apparently correlates with the living strategies of Mycobacteria − decreasing from free-living to opportunistic and strictly pathogenic species − while this is not true for CYP124 (Fig. 3). Additionally, the CYP125 gene is located within the cholesterol entry and catabolism cluster (Fig. S7), while CYP124 is located outside this cluster, suggesting its distinct transcriptional regulation. In addition, as described in the section “Structural basis for oxysterol hydroxylation”, CYP124 and CYP125 hydroxylate cholestenone to different stereoisomers, suggesting the involvement of the products in different metabolic pathways. We therefore conclude that despite similar ligand specificity, CYP124 and CYP125 have different functional roles in Mtb. This point is further supported by a consideration of genome sizes in the Mycobacterium genus (Fig. 3). Pathogenic bacteria often have smaller genomes than their non-pathogenic or less-pathogenic relatives − a phenomenon known as reductive genome evolution (*26*). This is in particular true for the Mycobacterium genus (see Fig. 3). Yet, both the CYP124 and the CYP125 genes are present in the whole MTC group, which additionally points to their different functional roles.

**Fig. 3.**
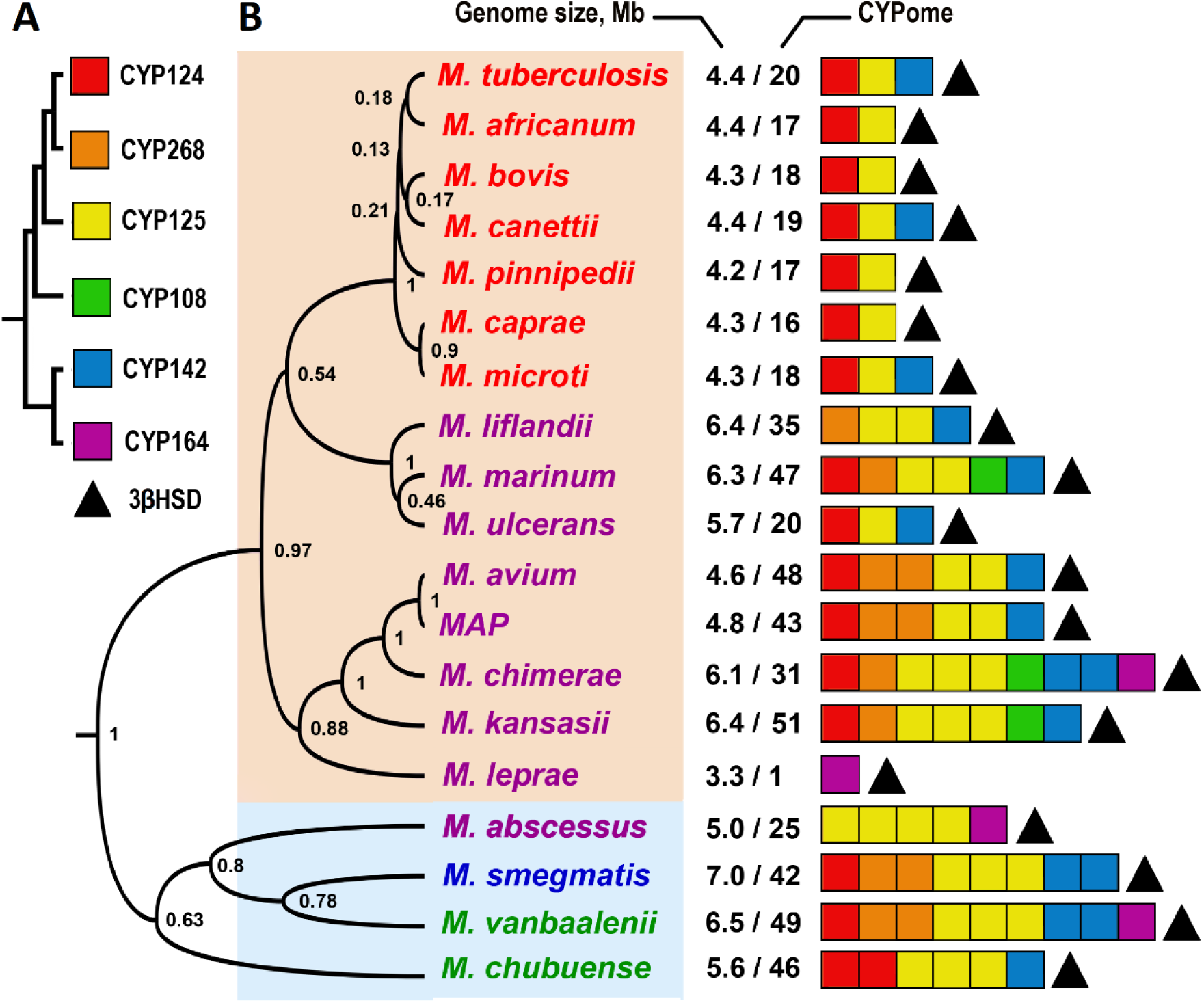
Distribution of steroid metabolizing CYPs and 3βHSD genes mapped onto a Bayesian inference phylogenetic tree based on 16S rRNA of the *Mycobacterium* genus. **A**. Phylogenetic relationships of CYPs (detailed in Fig. S6). **B**. Distribution of steroid metabolizing CYPs and 3βHSD across Mycobacteria species. The colored squares depict the CYP isoform, and the filled triangles depict the 3βHSD found in the listed species. Free-living non-pathogenic species are shown in green, commensal species in blue, species that cause diseases other than TB are in purple, and members of the MTC group are in red. Numbers at nodes indicate posterior probability. Slowly growing mycobacteria are shown on beige background, and fast growing species are shown on light blue background. The ratio between the genome size and the number of CYP genes is given in the middle.

We further analyzed co-regulation of steroid-metabolizing CYPs and 3βHSD in the Mtb transcriptome (*27*) with proteins engaged in transport of sterol derivatives into mycobacteria or known for their steroid-metabolizing function (Table S4). There are 27 transcriptional factors (TFs) of 206 analyzed that modulate the activity of steroid-metabolizing CYPs. The 3βHSD is translated in Mtb at the constitutive level throughout the entire life cycle. It is conservative across Mycobacteria (see Fig. 3). Therefore, 3βHSD is crucial for the Mtb life cycle. Meanwhile, the CYP124 and CYP125 genes are co-induced with TFs activated mainly on the earlier stages of infection and are coregulated with mce4 and mce3, correspondingly. It has been shown that ABC-like mce transporters supply Mycobacteria with hydrophobic molecules including sterols (*28*). Four mce operons have been identified in Mtb: mce1, mce2, mce3 and mce4 (*29*). The mce4 operon is involved in the transport of sterols (*30*). Therefore, we suggest that mce4 can deliver immunoactive steroid molecules to CYP124.

CYP125 is co-localized with mce4 in the cholesterol entry and catabolism gene cluster (Fig. S7) (*31*). However, CYP125 is not co-regulated with mce4, but it is strongly co-regulated with its paralogous transporter mce3 instead. The mce3 and mce4 transporters share a significant degree of similarity, thus raising a possibility that mce3 may also be involved in the translocation of steroid-like molecules. This functional interplay between CYP125 and mce4 within the cholesterol catabolism gene cluster from one side, and CYP124 and mce3 transporter from the other, broadens the regulatory network of the metabolism of sterols in Mtb that is active at the initial phase of infection.

A number of genes of orphan enzymes potentially involved in sterol metabolism are also co-regulated with CYP124 and CYP125 (Table S4). One of them is a putative 7α-hydroxysteroid dehydrogenase (7αHSD, gene Rv0927c), and it might be responsible for the dehydrogenation of a hydroxyl group at position C7 of the steroid skeleton (*32*). This enzyme belongs to a family of short-chain dehydrogenases/reductases (SDRs) that play a pivotal role in cell metabolism. The 7αHSD might possibly convert the inert for studied CYPs steroid 7α,25diOHC to the 7-ketosteroid, which in turn may be effectively metabolized by three steroid-metabolizing CYPs (Table 1). Another enzyme co-regulated with CYP125 is the 3α,20β-hydroxysteroid dehydrogenase fabG3 (Rv2002), which has an established role in steroid metabolism (*33*). In addition, a 3-ketosteroid Δ^1^-dehydrogenase kstD (Rv3537) is co-regulated with the studied CYPs and is predicted to be involved in A-ring degradation of 3-ketosteroids (*34*), which are putative products of 3βHSD.

This analysis highlights the complex regulatory network for uptake and diverse modifications of steroid-like molecules of the host.

### Inhibition of sterol-metabolizing Mtb CYPs

To test the effect of suppression of the enzymes involved in degradation of oxysterols, we selected imidazole derivatives. Azoles are known inhibitors of many CYPs, both human and bacterial/fungal, and they have potent anti-Mtb activity. We tested two imidazole derivatives: carbethoxyhexyl imidazole (CHImi, Fig. 4C) and carboxyheptyl imidazole (ChpImi), both containing an aliphatic chain crucial for binding to the selected CYPs. As expected, CHImi induced type II spectral response in all the studied CYPs (Fig. S1, S2, and S3 for CYP124, CYP125, and CYP142, respectively), this typically indicating direct coordination of a lone pair of electrons on the nitrogen of the imidazole ring with heme iron. In contrast, ChpImi shows substrate-like type I binding for CYP124 and CYP125, but not for CYP142, where a typical type II response is observed. Binding of ChpImi likely displaces a water molecule bound to the heme iron but prevents its direct coordination with iron. Alternatively, this can be explained by flipped orientation in the active site with the alkyl chain facing the heme. The two inhibitors show similar binding affinity to the studied CYPs (Table S2), while CHImi is more selective towards CYP124 and ChpImi is more selective towards CYP125.

**Fig. 4.**
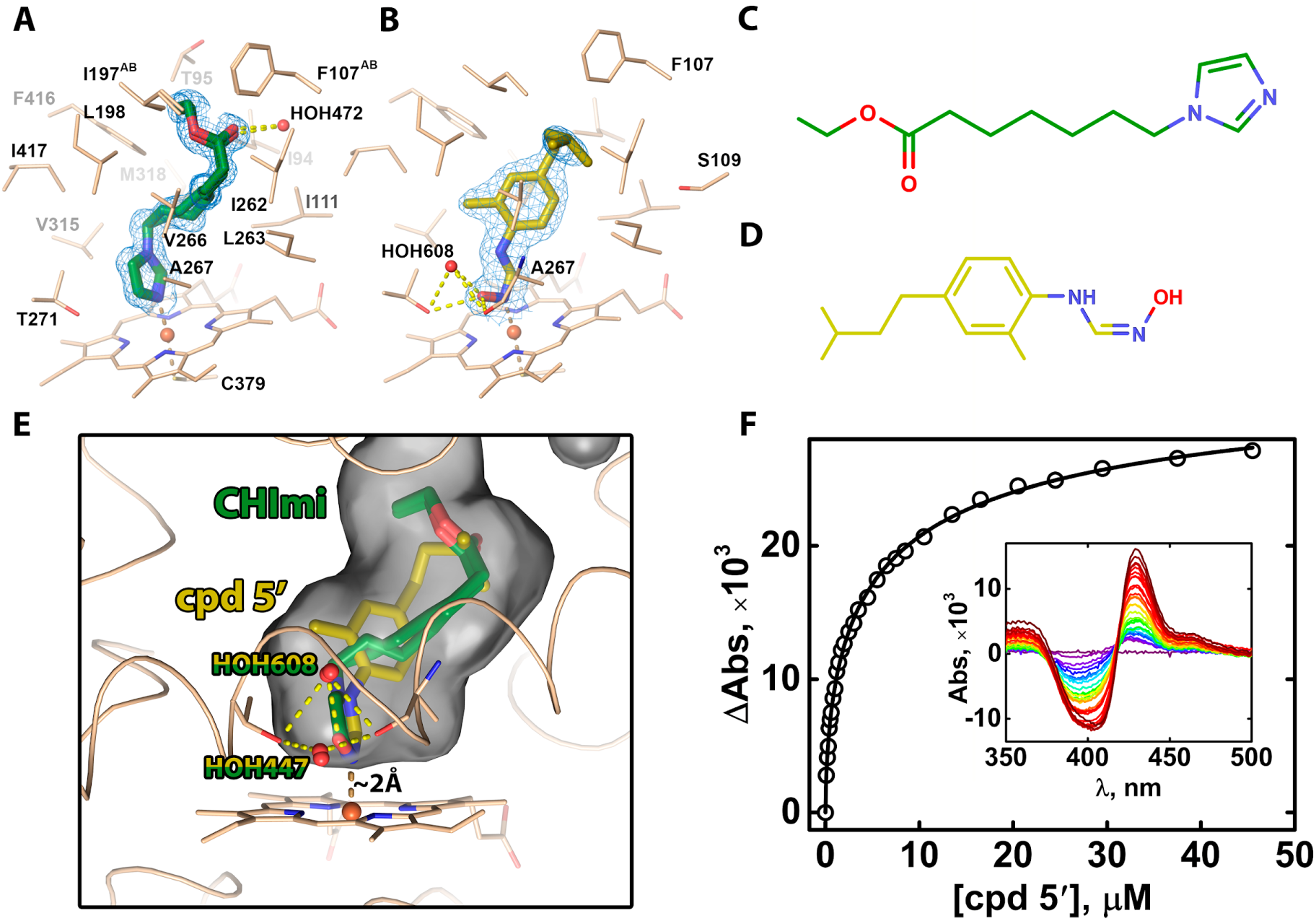
Structures of CYP124 in complexes with inhibitors. Structure of CYP124 binding pocket with inhibitors: CHImi, **A;** cpd5′, **B**. Only side chains within 5 Å and directly H-bonded water molecules or residues are shown for clarity. The *2mF*_*o*_−*DF*_*c*_ composite omit maps are contoured at 1σ. **C** and **D**, structures of CHImi and cpd5′. **E**, superposition of CYP124 inhibitors in the active site. **F**, CYP124 spectrophotometric titration with cpd5′ with difference spectra in inset.

We solved the structure of the CYP124–CHImi complex with 1.2 Å resolution (Table S3). CHImi has two very similar binding orientations in the main CYP124 cavity (Fig. 4A and Fig. S5D for electron density), both in the typical geometry for direct heme coordination. The binding site adopts a closed conformation with collapsed cavity I that is similar that of VD3/1αOHVD3 complexes (compare Fig. 2F and H/I).

Having the high-resolution structures in hand, we searched for an inhibitor with alternative scaffold with high affinity and specificity toward the studied CYPs. We designed a new phenylformamidamine derivative − N′-hydroxy-N-(4-isopentyl-2-methylphenyl)formimidamide (further referred to as cpd5’, see Supplemental Methods for synthesis details). It contains an oxime N-atom for coordination with heme and a short methyl-branched chain as in sterols (Fig. 4D). The compound shows high affinity to CYP124 as well as to CYP125 and CYP142 (Table S2, Fig. 4F).

We also performed inhibition assays (see Methods for details). Cpd5’ effectively inhibits hydroxylation of cholestenone by CYP124 with IC50 = 2.7 ± 0.5 μM). We further checked for its capacity to inhibit intracellular growth of Mtb in macrophages. For that, murine peritoneal macrophages were infected with Mtb H37Rv, and the fraction of macrophage-residing bacteria was estimated after incubation with or without cpd5’ (see Methods for details). The presence of cpd5’ reduces the number of macrophage-residing bacteria in a dose-dependent manner with IC50 = 18.2 µM (Table S5).

To visualize the mode of cpd5’ binding, we solved the structure of the CYP124–cpd5’ complex with 1.8 Å resolution (Table S3). The electron density fully describes the ligand (Fig. S5E). As expected, cpd5’ coordinates the heme iron with its oxime N-atom; an additional H-bond is formed between the oxime O-atom and HOH608 (Fig. 4B). The side-chain fits into the cavity, with its methyl group occupying the same place as in steroids (Fig. 4E).

Notably, in both CYP124−inhibitor complexes solved here (with CHImi and cpd5’), two water molecules (HOH447 and HOH608) enter the I-helix groove, disrupting its helical structure (Fig. 4E). This network resembles an oxygen-bound complex that is observed for P450cam and P450eryF, where two water molecules were suggested to transfer protons to the ferryl species during the catalytic cycle (*35, 36*). Thus, inhibitor binding might “trap” CYP124 in an intermediate state, which prompts the design of suicidal inhibitors or prodrugs.

## Discussion

Oxysterols are a re-emerging field of research because novel physiological activities of cholesterol derivatives are being uncovered that are beyond the regulation of cholesterol homeostasis. They are implicated in neurodegenerative diseases, the Hedgehog developmental pathway, and immune system function (*37*). Vitamin D derivatives, besides the vitamin’s classical role in calcium homeostasis, also modulate processes important for immunity (*38*). Here we explore at the molecular level the role of sterol derivatives, both oxysterols and secosterols, in host−pathogen interactions during Mtb infection.

We found that steroid-metabolizing enzymes can hydroxylate a number of immunoactive molecules in humans. Figure 1 summarizes our main findings in the context of related human immune and cholesterol-metabolism pathways. In humans, oxysterols are produced from cholesterol both enzymatically and/or non-enzymatically via free radical-mediated oxidation (blue background in the figure). Formation of 25OCH is catalyzed by cholesterol 25-hydroxylase (CH25H). 25OHC is a native ligand of LXR, which couples with retinoid X receptor (RXR) to promote transcription of the ABCA1, ABCG1, and ApoE genes (*39*). Products of these genes are involved in cholesterol transport and homeostasis as well as in regulation of the innate immune response (*40*). The LXRs modulate cell proliferation (*41*) and promote macrophage survival by inhibiting pro-inflammatory and pro-apoptotic factors during infection (*42*). 25OHC also has antiviral activity associated with alteration of the cholesterol content in late endosomes (*43*) and plasma membrane (*44*), thus impairing viral particle assembly and fusion with membranes (*13*). Similar reduction in plasma-membrane-accessible cholesterol induced by 25OHC was recently demonstrated for bacterial infections (*45*). Our findings suggest that the Mtb enzymatic system can modulate the activity of 25OHC, enabling it to interfere with the human immune response. This is consistent with increased mortality from Mtb coinfection with influenza and other viral infections (*46, 47*), which may be attributed to the decreased antiviral effect mediated by 25OHC.

How Mtb can access 25OHC during infection is not yet explored. It might be associated with mce proteins − lipid/sterol transporters that are suggested to modulate host cell signaling. In particular, mce4 has a high affinity to 25OHC (*48*), and it is upregulated along with CYP124 as suggested by our bioinformatic transcriptome analysis (Table S4).

7α,25diOHC can be formed from 25OHC by CYP7B1 (*49*). 7α,25diOHC is a very potent agonist of the EBI2 receptor, which is involved in regulation of chemotactic migration of B-cells and dendritic cells (*50, 51*). Its level is strictly regulated by 3β-hydroxysteroid dehydrogenase type 7 (*13*). Our data suggest that mycobacterial 3βHSD might functionally mimic this human enzyme, interfering with oxysterol-mediated migration of immune cells. The function of 3βHSD in immune response may be associated with the increased number of granulomas in the lungs of animals infected by a 3βHSD mutant strain (*6*). The absence of 3βHSD in Mtb may result in a more activated EBI2 pathway, hence more pronounced immune response and, as consequence, granuloma formation. 7α,25diOHC may also be transformed to its keto form by a putative 7αHSD identified in our transcriptome analysis. 7keto,25OHC can further be oxidized by CYP124/CYP125 and to a lesser extent by CYP142.

In the vitamin D activation pathway (Fig.1, green), biologically inert VD3 is hydroxylated to 25OHVD3 by CYP2R1 in the human liver and further to an active hormonal form, 1α,25diOHVD3 or calcitriol, by CYP27B1 in the kidney (*52*). Synthetically-derived 1αOHVD3 (alfacalcidol (*53*)) can also be utilized as a 1α,25diOHVD3 precursor. 1α,25diOHVD3 binds to vitamin D receptor (VDR) and induces transcription of antimicrobial peptides – cathelicidin and β-defensin 2 (*14*). The most potent ligand, 1α,25diOHVD3, is not metabolized by the studied enzymes. However, its precursors – VD3 and 1αOHVD3 – are hydroxylated by CYP124. The ability of CYP124 to metabolize VD3 and its derivatives is consistent with the observation that high doses of VD3 supplementation is not efficient for anti-TB therapy (*54*).

1α,20S-dihydroxyvitamin D3 (1α,20S-diOHVD3) is a natural VD3 metabolite that interacts with VDR with potency similar to that of 1α,25diOHVD3 (*55*). It is structurally similar to 1αOHVD3 and 20SOHC, and both are CYP124 substrates (Table 1). Even though we did not study 1α,20S-diOHVD3 experimentally here, we expect it to be a CYP124 substrate as well.

7DHC is a precursor for the most diverse set of oxysterol products that have been observed to date (*16*). It links together two classes of steroids studied in our work, since it is the common biosynthetic precursor of cholesterol and VD3 (shown on yellow background in Fig. 1). Notably, in our study 7DHC was among the most actively metabolized sterols by CYP124 and CYP125 (Table 1). These reactions may put out of action a wide range of 7DHC derivatives, producing massive modification of related host pathways.

7DHC can be converted by human CH25H to 25OH7DHC − an agonist for both LXRs and VDR discussed above as targets for 25OHC and 1α,25diOHVD3, respectively (*56*). CYP124 metabolizes 25OH7DHC with efficiency similar to that of 25OHC.

Altogether, evolutionarily conserved Mtb enzymes, constitutively expressed 3βHSD and CYPs induced at the initial stages of infection, metabolize different immune oxysterols and secosteroids, thus interfering with a broad range of signaling pathways of the host. The bioinformatic, functional, and structural data provided here could potentially open a next generation of pharmacological approaches aimed at breaking pernicious interrelations between Mtb and the human immune system. The structural basis established by our study has the potential to defeat the most severe cases of multi-drug-resistant TB.

## Methods

### Cloning, expression, and purification of recombinant proteins

cDNAs encoding the 3βHSD (gene Rv1106c) and CYP125 (gene Rv3545c) and CYP142 (gene Rv3518c) and CYP124 (gene Rv2266) were amplified by PCR of genomic DNA of Mtb H37Rv (kindly provided by the Vyshelessky Institute of Experimental Veterinary Medicine, Belarus). Expression plasmids for each protein were generated using vector pTrc99a. The proteins were expressed and purified as described previously (*12*). The cDNA encoding spinach Ferredoxin-1 (Fdx-1) (used in catalytic activity assay) was amplified from the total RNA isolated from *Spinacia oleracea* seedlings. Fdx-1 was expressed in *E. coli* and purified using metal-affinity and anion-exchange chromatography. Ferredoxin reductase Arh1 (mutant A18G) was provided by Prof. Rita Bernhardt (Saarland University, Saarbrucken, Germany), expressed and purified as described in (*57*).

### Ligand binding studies

To determine ligand-binding constants (Kd_app_ values) of the CYPs, spectrophotometric titration was performed using a Cary 5000 UV-Vis NIR dual-beam spectrophotometer (Agilent Technologies, Santa Clara, CA) in 1 cm quartz cuvettes. Stock solutions of the steroids were prepared at concentration 10 mM in 45% hydroxypropyl-beta-cyclodextrin (HPCD). The titration was repeated at least three times, and Kd_app_ was calculated as described previously (*12*).

### Catalytic activity assay

Catalytic activity of the Mtb CYPs was reconstituted in 50 mM potassium phosphate (pH 7.4) containing 0.5 µM CYP, 2 µM spinach Fdx1, 0.5 µM Arh1, 100 µM substrate, 1 mM glucose 6-phosphate, 1 U/ml glucose 6-phosphate dehydrogenase, and 0.4 mM β-NADPH. The proteins (CYPs, Fdx1, and Arh1) were pre-incubated with the steroid substrates in the buffer solution for 10 min at 30°C, and the reaction was started by adding the NADPH-regenerating system containing glucose 6-phosphate, glucose 6-phosphate dehydrogenase, and β-NADPH. After 2, 5, and 30 min of incubation at 30°C, aliquots of the reaction mixture were taken; the enzymatic reaction was stopped by boiling. The extracted with methylene chloride lipids were subjected to HPLC analysis. HPLC with DAD was performed on an Agilent 1200 Series instrument equipped with an Eclipse Plus C18 column (4.6 × 250 m; particle size = 5 μm; Agilent). Activities of the enzymes were estimated as Turnover Number nmoles of metabolized product / nmol of CYP / min (min^-1^). The site of hydroxylation of sterols was estimated using the Agilent HPLC 1200 instrument equipped with an Agilent Triple Quad 6410 mass detector. The chromatographic analysis was carried out on a Zorbax Eclipse XDB C18 column (4.6 × 150 mm; 5.0 µm).

To evaluate the inhibitory effect of cpd5’ on the cholestenone hydroxylation activity of CYP124, cpd5’ was added with substrate to the final concentration, ranging from 0.1 to 50 µM. The reaction was stopped after 30 min, and the reaction mixture was extracted and subjected to HPLC analysis as described above.

Catalytic activity of the mycobacterial 3βHSD was determined in a reconstituted system containing the following components: 100 mM Tris-HCl buffer (pH 8.5), with 150 mM NaCl, and 30 mM MgCl_2_, 1 µM of the 3βHSD, 100 µM progesterone as internal standard, and 100 µM of the substrate (10 mM stock solution in 45% HPCD). The reaction was initiated by addition of 2.8 mM β-NAD. After 30 min of incubation at 30°C, the reaction was stopped by addition of methylene chloride, and the reaction mixture was extracted and subjected to HPLC analysis.

### NMR analyses

^1^H NMR (499.93 MHz) and ^13^C NMR (125.72 MHz) spectra were recorded on a Bruker AVANCE-500 NMR spectrometer. CDCl_3_ was used as the solvent, and the residual solvent signals (δ 7.26 ppm for ^1^H NMR and 77.16 ppm for ^13^C NMR) served as an internal reference standard. COSY, HSQC, and HMBC experiments were carried using the standard Bruker software.

### Analysis of metabolism of 25OHC by Mtb cells

Mtb H37Rv was cultured for 8 days at 37 °C, 5% CO_2_ in 24 well plate in Sauton medium with 0.025% Tween 80 at 10^6^ Mtb/ml/well. 25OHC was added to sample wells at the final concentration 10 µM. The calculated Mtb cell content was 3.5 ± 0.4 × 10^7^ CFU H37Rv/well. After 0, 6 or 24 hours of incubation, sodium azide (1% final concentration) was added and the samples were incubated at 96 °C for 1 min. The steroid fraction was extracted and subjected to LC-MS analysis. LC-MS experiments were carried out with the Agilent HPLC 1200 instrument equipped with the Agilent Triple Quad 6410 mass detector. The chromatographic analysis was carried out using a Zorbax Eclipse XDB C18 column (4.6 × 150 mm; 5.0 µm). The mobile phase consisted of two solvents: (A) 0.1% (v/v) formic acid in water, and (B) 0.1% (v/v) formic acid in methanol. The linear gradient started with 85% B and reached 100% B at 5 min. The flowrate was 0.6 ml/min, and the injection volume was 10 μl. Mass spectrometry experiments were performed with an electrospray ionization (ESI) source in positive-ion mode. The nebulizing gas flow rate was set at 10 l/min, the fragmentor at 90 V, drying gas temperature at 350 °C, capillary voltage at 4000 V, and nebulizer at 30 psi. The data acquisition mode was MS2Scan from 280 to 500 Da. Metabolic activity of the Mtb cells was estimated by consumption of the 25OHC substrate.

### Crystallization, data collection, and crystal structure determination

CYP124 was crystallized using a sitting drop approach in 96-well crystallization plates with commercially available kits (Qiagen) at 20 °C with 1:1 protein/mother liquor ratio with the ligand concentration of 100 μM. The CYP124(A65T mutant)–cholestenone complex was crystallized in 0.1 M MOPS pH 6.5, 27% PEG 3350; CYP124–VD3, CYP124–1αOHVD3, and CYP124–CHImi complexes were crystallized in 0.2 M Mg(Cl)_2_, 0.1 M Bis-Tris pH 6.5, and 25 % PEG 3350. The same conditions but without the salt worked for the CYP124–cpd5’ complex. Glycerol (20%) as cryoprotectant was added before flash-freezing under liquid nitrogen when necessary.

Diffraction data were collected at the European Synchrotron Radiation Facility (ESRF) beamlines id23-1 (for the CYP124–cholestenone complex), id30b (for the CYP124–VD3, CYP124–1αOHVD3, and CYP124–CHImi complexes), and id30a1 (for CYP124–cpd5’). The data collection strategy was optimized in BEST (*58*). All data were processed in the XDS software package (*59*). Crystallographic data collection statistics are given in Table S3.

The phase problem was initially solved for the CYP124–cholestenone complex by molecular replacement in PHENIX.Phaser (*60*) with the poly-ala model generated from substrate-free CYP124 (PDB ID code 2WM5). The poly-ala model of the refined CYP124–cholestenone complex was later used as a search model for all the other CYP124 complexes.

In all cases except for cpd5’, the space group was P12_1_1 and contained four molecules per asymmetric unit for CYP124–cholestenone and one molecule per asymmetric unit for all the others. For the cpd5’ complex, the space group was P4_3_2_1_2 with one molecule per cell.

The model was built in PHENIX AutoBuild (*61*). PHENIX.Refine (*62*) was used for refinement. COOT (*63*) was used for manual refinement. On the last refinement steps, the *mF*_*o*_−*DF*_*c*_ map unambiguously showed the ligands (Fig. S5). Final resolution cut-off was determined by application of paired refinement (*64*). The quality of the resulting model was analyzed by PHENIX. MolProbity (*65*) and the Quality Control Check server (https://smb.slac.stanford.edu/jcsg/QC/).

PHENIX.eLBOW (*66*) was used to generate restraints for 1αOHVD3 and CHImi, and cpd5’ restraints were generated in AceDRG (*67, 68*). The figures containing electron density and molecular structures were generated using PyMOL (*69*) and Chimera (*70*).

### Retrieval of P450 protein sequences, multiple sequence alignment, and phylogenetic tree construction

To build a set of P450s, we performed BLASTP and TBLASTN searches against the GenBank database using the sequences of three steroid-binding CYPs of Mtb H37Rv (CYP124, CYP125, and CYP142) as queries. Only those sequences that had more than 40% identity and E-value less than 10^− 7^ were included in our set. Accession numbers of sequences used are detailed in Fig. S7. The resulting sequence sets were aligned using MUSCLE (*71*). The datasets were analyzed using Bayesian inference implemented on BEAST version 2.0 (*72*). For phylogenetic analysis of the P450s, the JTT model with a discrete gamma (G) distribution across sites (four categories) was considered as the best fitting one, selected by MEGA software version 7.0 (*73*). The uncorrelated relaxed log normal clock model and a Yule birth model with birth rate gamma distribution (α = 0.001, β = 1000) were chosen to allow every branch of a tree to evolve with its own rate. Branch length in the resulting tree is proportional to the number of amino acid substitutions per site. The analysis was performed by running 10^7^ generations in two chains, keeping every 1000th tree. The results of the runs were summed with burn-in percentage 20 in TreeAnnotator and the consensus tree was created with the same burn-in percentage. The final tree was edited in FigTree.

### Reconstruction of *Mycobacterium* phylogeny and detection of distributions of P450s among species

To reveal the evolutionary history of the mycobacterium genus, the 16S rRNAs of 19 bacterial species were obtained with a BLASTN search against the RefSeq database using 16S rRNA of Mtb H37Rv as the query. The MUSCLE package was used to align these sequences, which was followed by phylogenetic analysis in BEAST 2.0. The alignment was analyzed with 10^7^ generations of MCMC under the HKY substitution model and with gamma distributed rates (5 categories), uncorrelated relaxed log normal clock model, and a Yule birth model with birth rate gamma distribution (α = 0.001, β = 1000). The consensus tree was created with burn-in percentage 20 in TreeAnnotator and edited in FigTree.

### Anti-Mtb activity

To evaluate the activity of cpd5’ against intracellular mycobacteria, peritoneal macrophages from C57BL/6 mice were cultured in RPMI-1640 medium supplemented with 2 mM L-glutamine, 10 mM HEPES, and 2% heat-inactivated FCS. The cells were seeded on 96-well plates at 60 × 10^3^ cells per well, incubated at 37 °C, 5% CO2 for 60 min, and infected with Mtb strain H37Rv Pasteur at MOI = 5. After 24 hours, extracellular mycobacteria were washed out twice, and cpd5’ was added to the infected macrophages for another 24 hours. The supernatants were aspirated, and the cells were washed out and lysed with H2O (0.1 ml per well) for 10 min at room temperature. After restoring ionic balance with 0.1 ml of 2 × Dubos broth mycobacteria released from macrophages were incubated for 3 days at 37 °C, 5% CO_2_. For the last 18 h of incubation, [^3^H]-uracil was added at 2 µCi per well. Incorporated radioactivity was estimated by transferring the content of the wells onto fiberglass filters and assessed using a Wallac 1409 scintillation counter.

## Data availability

Structures for CYP124 in complex with cholestenone, VD3, 1αOHVD3, CHImi, and cpd5’ have been deposited in Protein Data Bank with accession numbers 6T0F, 6T0G, 6T0H, 6T0K, and 6T0L, respectively. The diffraction images and processing data were deposited into the Integrated Resource for Reproducibility in Macromolecular Crystallography (http://proteindiffraction.org/). Other data and materials are available upon request from the corresponding authors.

## Acknowledgments

We thank S. Fatykhava and O. Bokut for excellent technical assistance. We thank Prof. Rita Bernhardt (Saarland University, Saarbrucken, Germany) for providing the expression construct for Arh1. This work was supported by a joint research grant of the Belarusian Republican Foundation for Fundamental Research (X18P-098) and the Russian Foundation for Basic Research (18-54-00030). Mtb whole-cell assays in part were done under the theme of the Central Research Institute for Tuberculosis 0515-2014-0055 “Efficacy of TB drug therapy in dependence of the host genetics”. This work was supported in part by a grant from the NIAID (F.A. OISE-9531011 – Identification of Molecular Targets for Anti-Tuberculosis Drug Discovery). Any opinions, findings, and conclusions or recommendations expressed in this work are those of the author(s) and do not necessarily reflect the views of NIAID. We acknowledge the ESRF Structural Biology Group and especially A. N. Popov for assistance with crystallographic data collection. Thanks to the NIH Joint Center for Structural Genomics for the structure quality validation server. We appreciate a kind gift of G. Marchale (Institut Pasteur, Paris, France), who provided us with Mtb H37Rv (Pasteur). We are grateful to Richard Lozier for editing our manuscript.

## Author’s contributions

Conceptualization, A.Gi., V.B., N.S.;

Methodology, A.R., A.M., V.B., A.Gi., N.S., A.B., R.L., A.H., D.H., A.A.;

Investigation, T.V, S.B., I.G, E.M., Ant.K, A.Gu., K.K., I.M., A.L., D.Z., R.A., M.S., S.S, Ann.K,

P.S, A.B, V.D, Y.C, A.S, R.L, A.H., E.S, A.R., A.M., A.Gi, D.E.H., B.N., K.M., A.A., A.R., N.S.;

Writing – Original Draft, N.S.;

Writing – Review & Editing, S.B., Ann.K, V.G., V.B., N.S, A.Gi.;

Funding Acquisition, V.G., V.B., N.S., A.Gi., Supervision, V.B., N.S., A.Gi.

## Declaration of Interests

A.Gi. and E.S. are employees of MT-Medicals LLC. The other authors declare no competing interests.

