## Supplemental Information for "Metabolic fate of human immunoactive sterols in *Mycobacterium tuberculosis*"

### Supplementary information

#### Fig. S1. Difference spectra and titration curves of Mtb CYP124 with various ligands

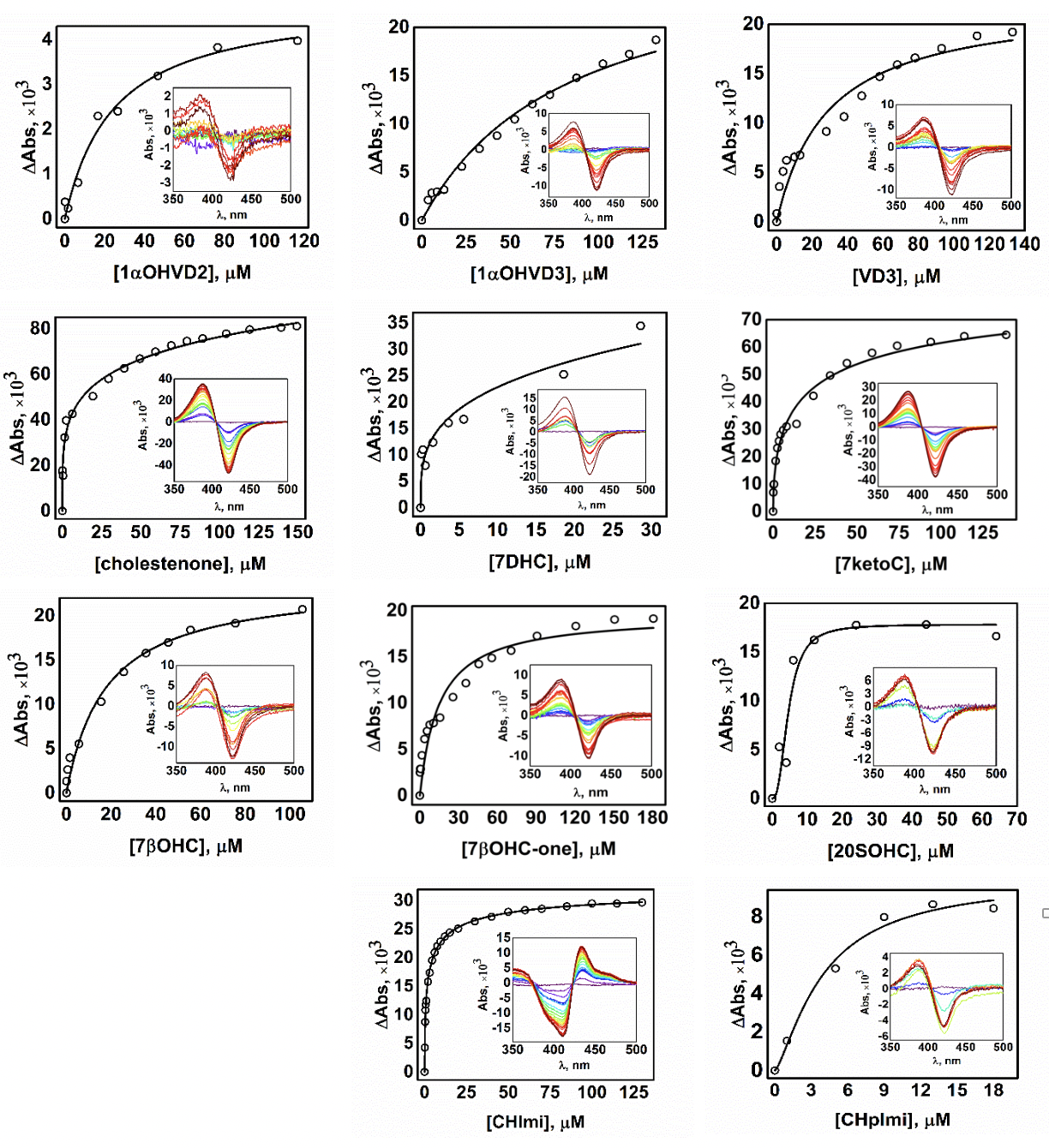

The concentration dependence of ligand binding was deduced from the difference absorption changes obtained from the titration of CYP124 (1-2 µM) with increasing concentrations of the ligands. Kd_app_ values are given in Table S2.

#### Fig. S2. Difference spectra and titration curves of Mtb CYP125 with various ligands

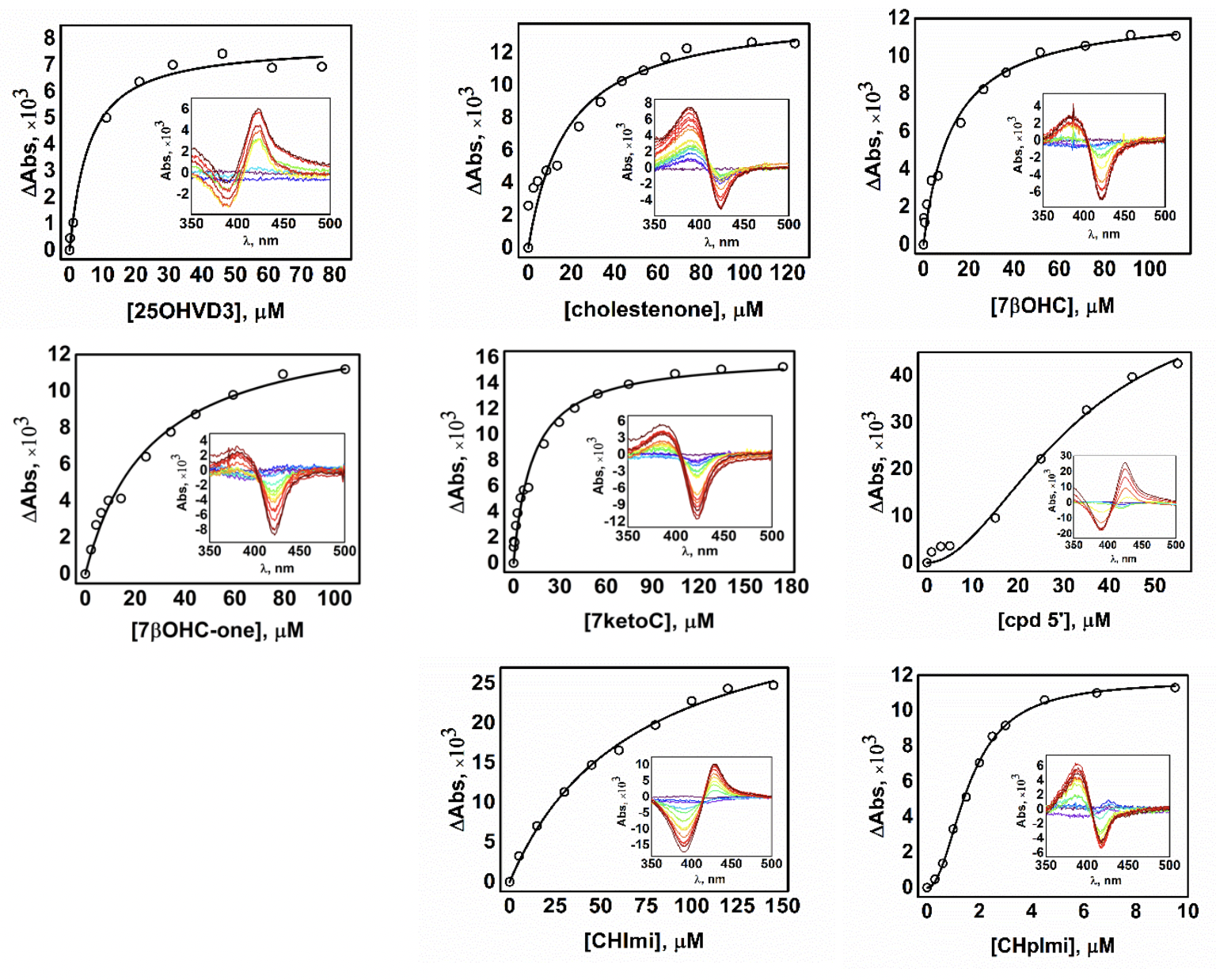

Concentration dependence of ligand binding deduced from the difference absorption changes obtained from the titration of CYP125 (1-2 µM) with increasing concentrations of the ligands. Kd_app_ values are given in Table S2.

#### Fig. S3. Difference spectra and titration curves of Mtb CYP142 with various ligands.

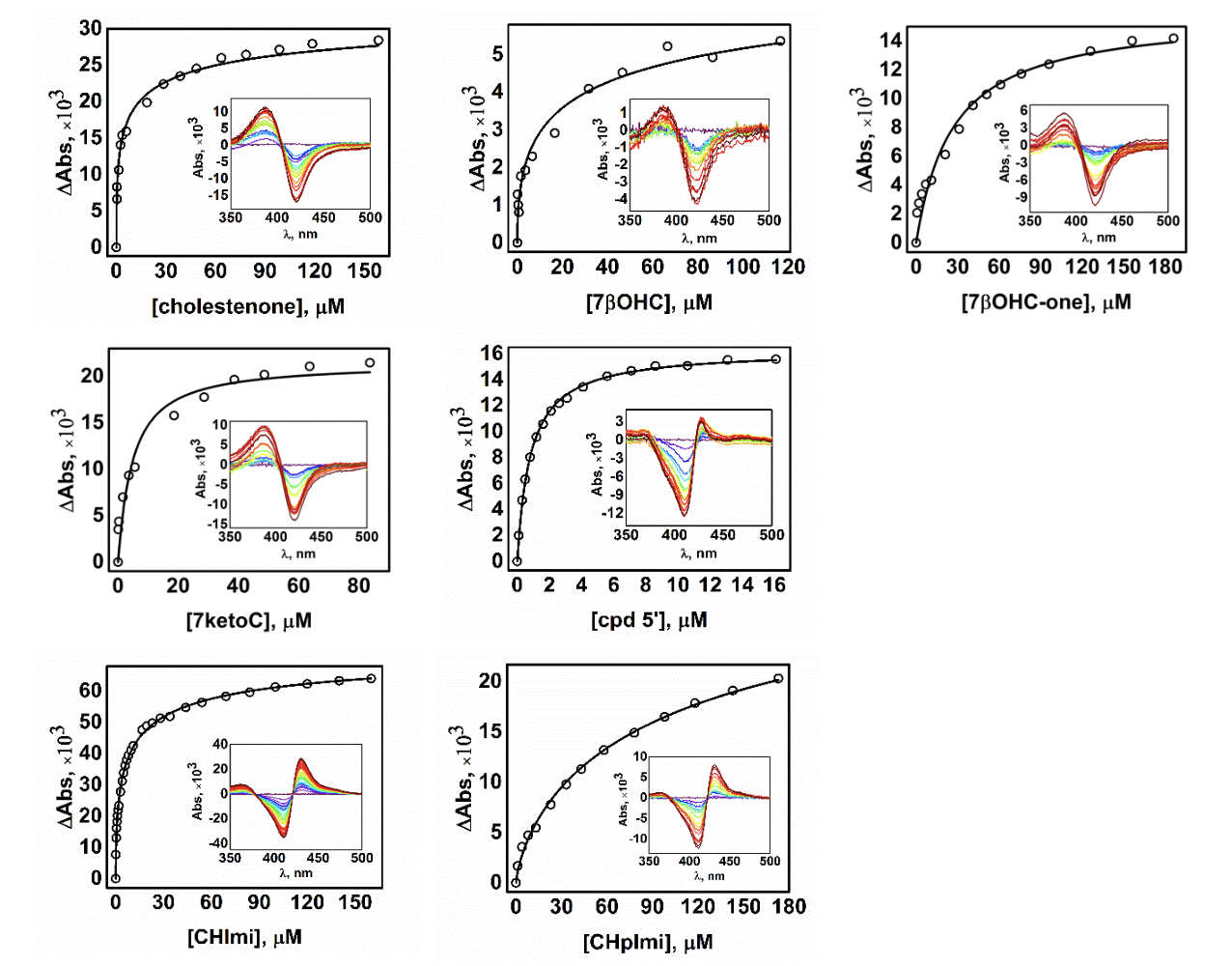

Concentration dependence of ligand binding deduced from difference absorption changes obtained from the titration of CYP142 (1-2 µM) with increasing concentrations of the ligands. Kd_app_ values are given in Table S2.

#### Fig. S4. NMR spectra and signal assignment of the hydroxylation product of 25OHC-one by CYP124 (25,26diOHC-one).

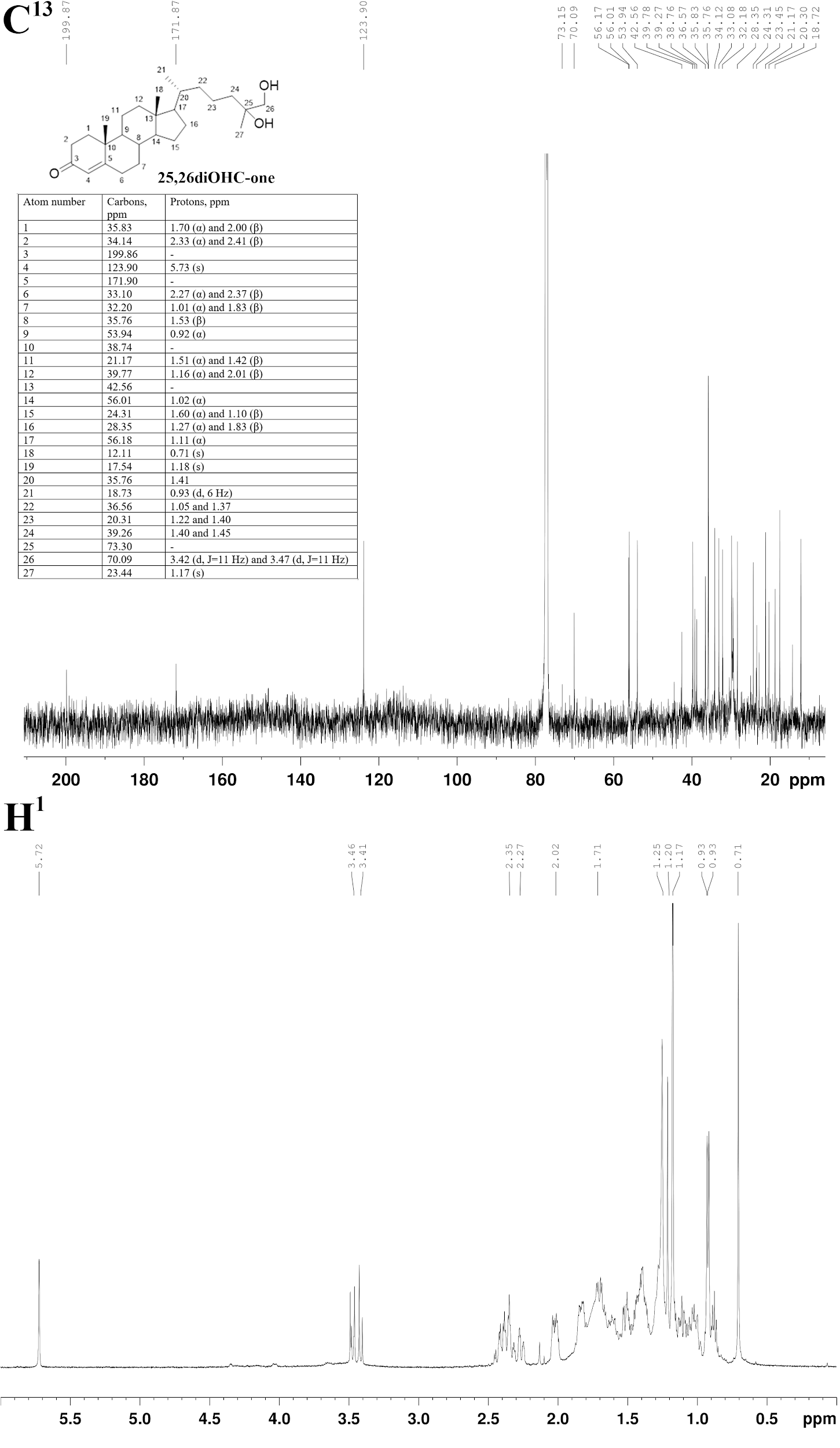

The isolated (after solid phase extraction) mixture of products analyzed by NMR correlation spectroscopy methods. The mixture was shown to contain two major components −25OHC-one (26/27 methyl groups at 1.20 ppm) and 25,26diOHC-one – at in approximately 1:4 ratio. Protons and carbon nuclei of the 25,26diOHC-one were assigned from the NMR spectra using the HSQC, COSY, and HMBC methods. The signals of methylene groups of the side chain were heavily overlapped; therefore, their assignment was done in analogy with known data for vitamin D derivatives (*74*). Satisfactory data of NOESY experiments could not be obtained due to insufficient amounts of 25,26diOHC-one. The configuration of chiral centers of the side chain remains unclear (*75*), and that of the steroid core are based on analogy with steroidal 4-en-3-ones (*76*, *77*).

#### Fig. S5. Ligand electron density maps of CYP124–ligand complexes

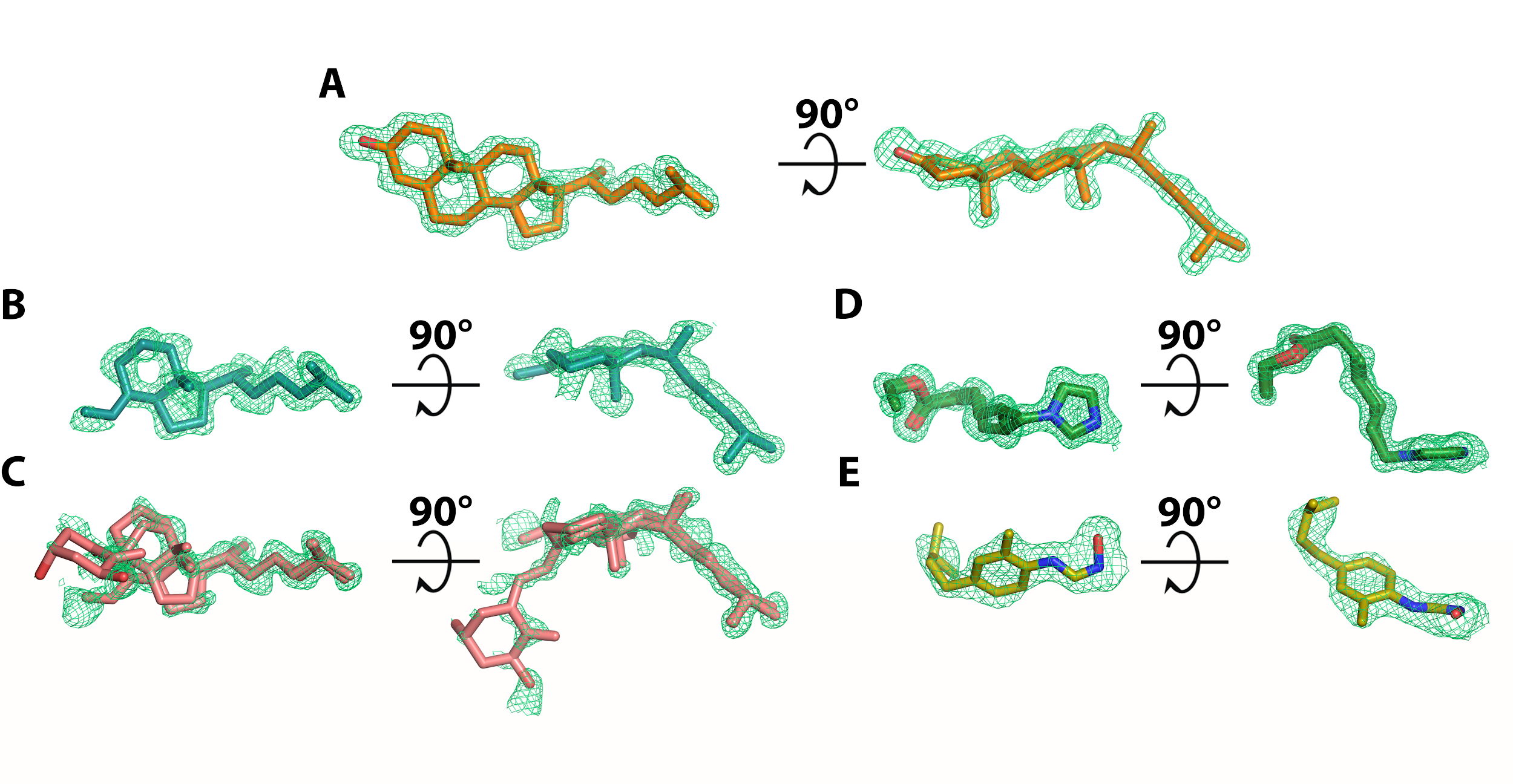

The *mF_o_−DF_c_* maps of CYP124 ligands: cholestenone, **A;** VD3, **B;** 1αOHVD3, **C;** CHImi, **D;** and cpd5', **E**. All of the difference maps are contoured at 3σ, except for 1αOHVD3, which is contoured at 2σ.

#### Fig. S6. CYPs phylogenetic tree of Mycobacteria constructed with Bayesian inference analysis

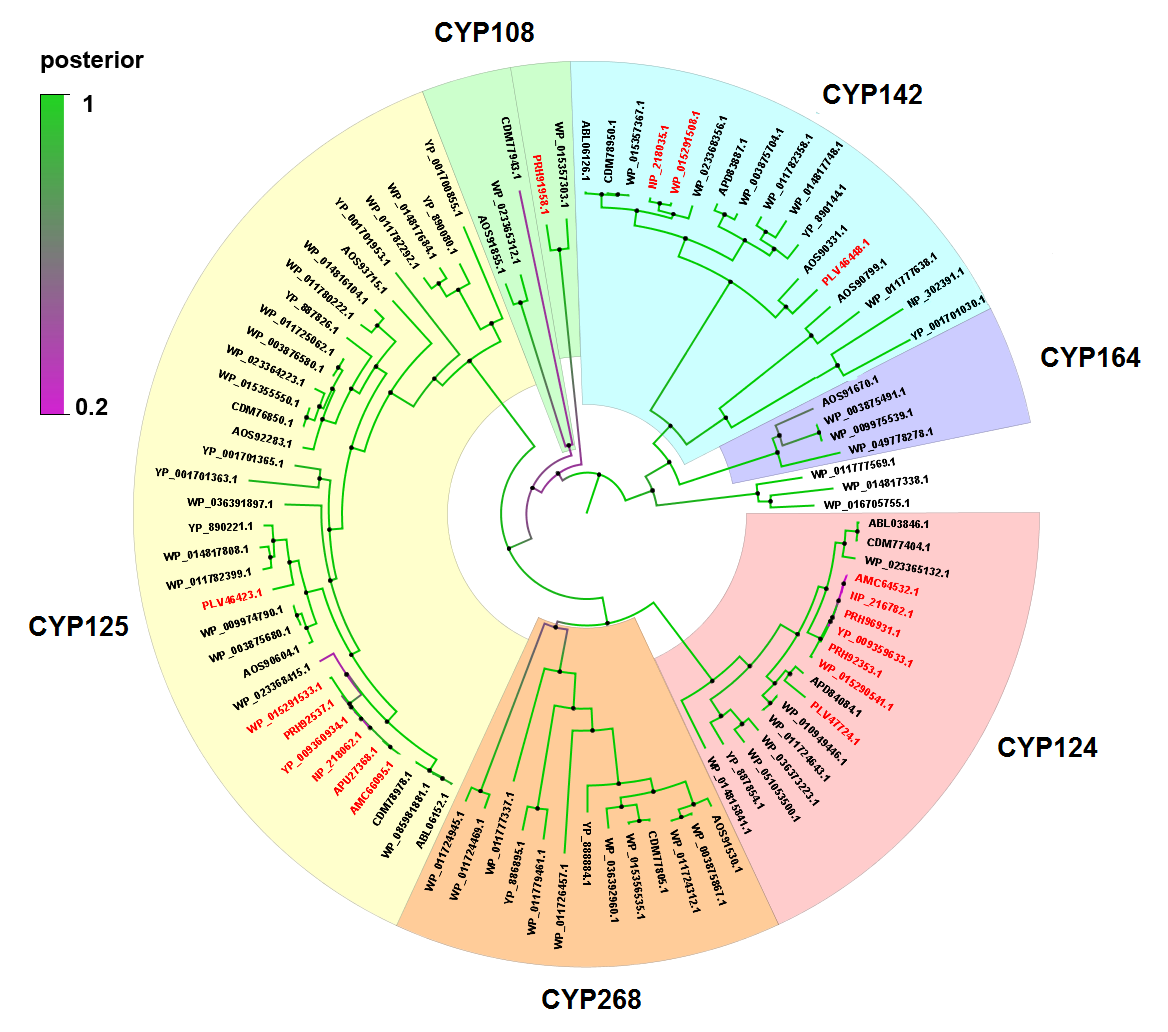

Members of the MTC group are shown in red. The colour of segments indicates posterior probability. The CYP142 clade was the first clade of steroid-metabolizing CYPs that diverged and is widely distributed in Mycobacteria compared with its paralog CYP164, which metabolizes fatty acids in *Mycobacterium smegmatis* (*78*). Using Mtb CYP124 as an input, sequence search among mycobacterial species with different living strategies revealed additional CYP families: fatty acid hydroxylase CYP268 and α-terpineol hydroxylase CYP108. The CYP268 family is paralogous to the CYP124 family, but it is not as widely distributed as CYP124.

#### Fig. S7. Number of homologs in the Mtb H37Rv genome mapped onto the cholesterol entry and catabolism cluster Rv3492c-Rv3575c

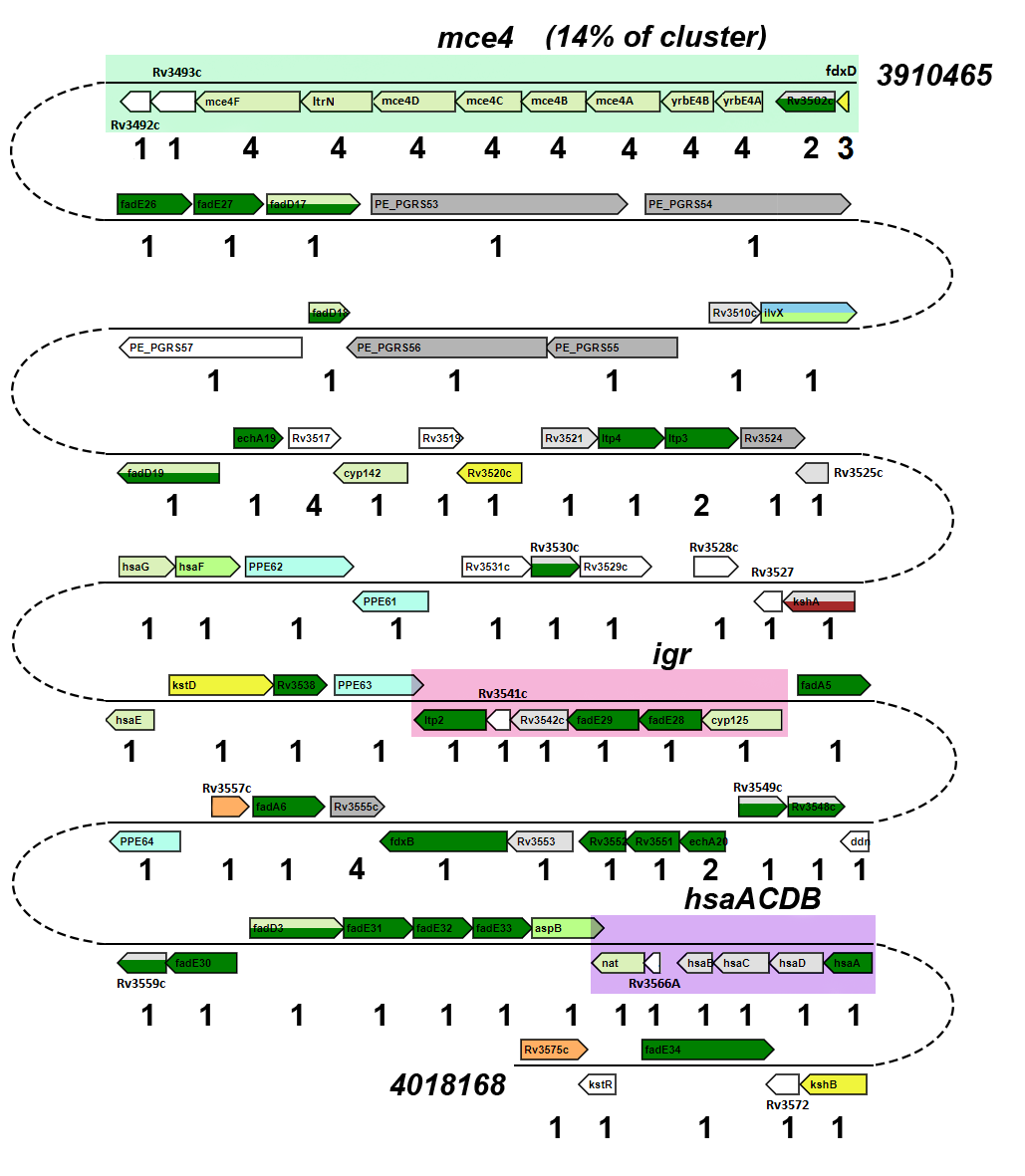

Annotated operons within the cluster are marked with colors. Only homologs located outside the cluster were considered. Genes were considered paralogs when they shared more than 30% identity. The visualization was made using the microbial genomic context viewer (MGcV) (*79*). The mammalian cell entry operon mce4 imports cholesterol for long-term survival of the bacilli (*80*). The intracellular growth operon (igr) codes for enzymes involved in β-oxidation of steroids (*81*). Operon hsaACDB is involved in cholesterol sterol‐ring degradation (*31*). Genes are colored according to their COG annotation: light green (yrbE4A-mce4F, cyp142, hsaG, hsaE, cyp125, nat) for secondary metabolite biosynthesis, transport, and catabolism; dark green (Rv3502c, fadE26, fadE27, fadD17, fadD18, fadD19, echA19, ltp4, ltp3, Rv3530c, Rv3538, ltp2, fadE29, fadE28, fadA5, Rv3548c, Rv3549c, echA20-Rv3552, fdxB, fadA6, Rv3559c-fadE33, hsaA, fadE34) for lipid transport and metabolism; bright green (ilvX, hsaF, aspB) for amino acid transport and metabolism; yellow (fdxD, Rv3520c, kstD, kshB) for energy production and conversion; orange (Rv3557c, Rv3575c) for transcription; light blue (PPE62, PPE61, PPE63, PPE64) for cell motility; bright blue (ilvX) for coenzyme transport and metabolism; red (kshA) for inorganic ion transport and metabolism; gray (PE_PGRS53, PE_PGRS54, PE_PGRS55, PE_PGRS56, Rv3524, Rv3555c) for proteins with unknown function.

#### Fig. S8. Scheme of synthesis of 25OH7DHC and cpd5'

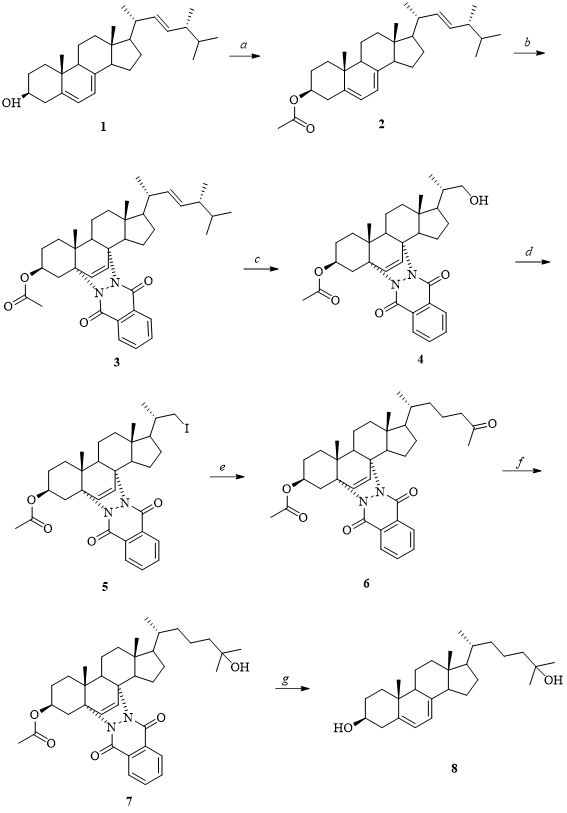

**1a**: Ac_2_O, Py, 70%; **b:** Phthalic hydrazide, Pb(OAc)_4_, CH_2_Cl_2_, 0℃, 94%; **c:** O_3_, CH_2_Cl_2_/MeOH, –60℃, then NaBH_4_, 63%; **d:** I2, ImH, PPh3, CH_2_Cl_2_, 93%; **e:** 3-buten-2-one, Zn-Cu, NiCl_2_, Py, 76%; **f:** MeMgI, Et_2_O, then NH_4_Cl, 48%; **g:** LiAlH_4_, THF, 66℃, 79%. See Supplementary Methods for details.

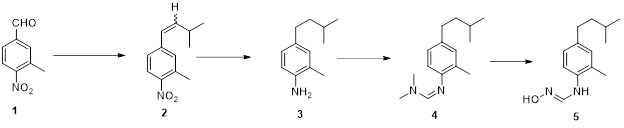

**1′	2′	3′	4′	5′**

(E)-N'-hydroxy-N-(4-isopentyl-2-methylphenyl)formimidamide (compound 5', cpd5') was prepared from 3-methyl-4-nitrobenzaldehyde in five steps. The synthesis includes the Wittig reaction, catalytic reduction of nitro- and double bonds over Raney nickel, the treatment of the resulting aniline with N,N-dimethylformamide dimethyl acetal followed by hydroxylamine hydrochloride. See Supplementary Methods for details.

#### Table S1. Biosynthesis and effects of oxysterols and VD3 derivatives on immune cells

|  | **Structure** | **Enzyme** | **Known Receptor** | **Function** |
| --- | --- | --- | --- | --- |
| 1 | 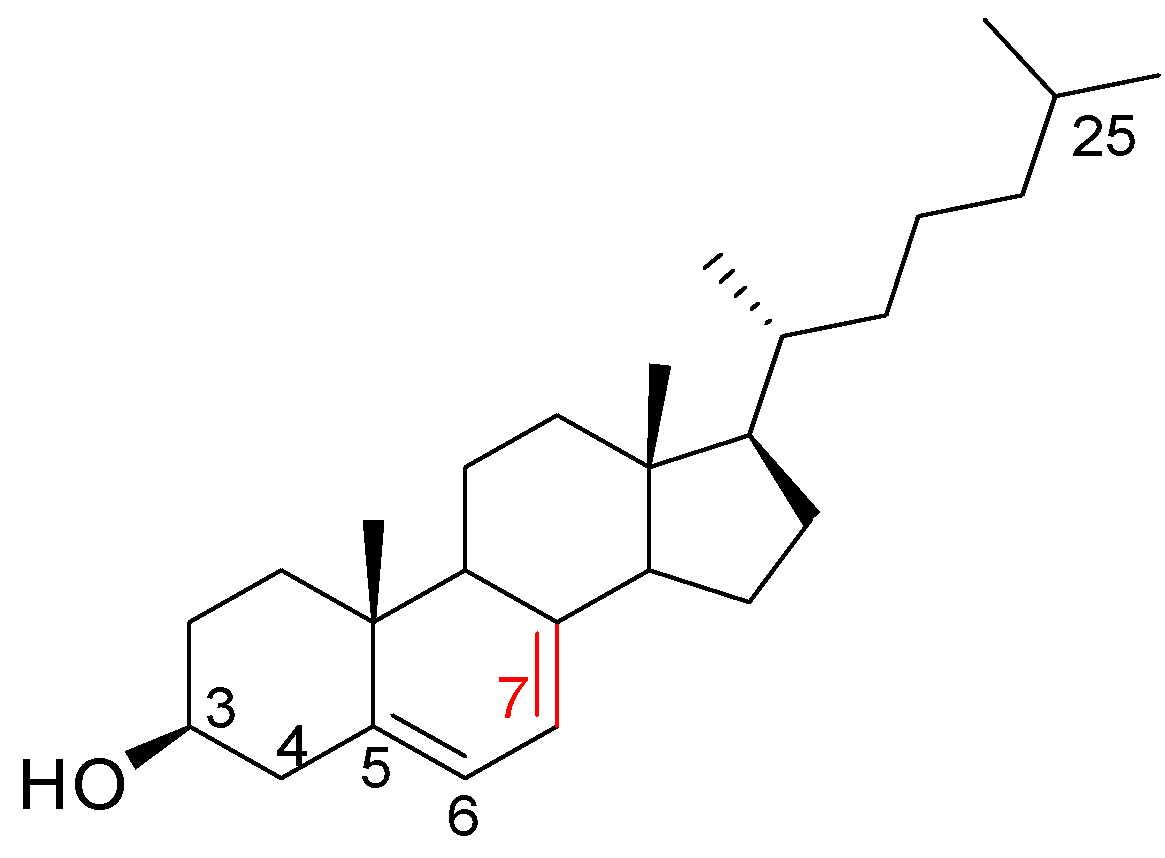  7-dehydrocholesterol  (7DHC, provitamin D3) | Sterol ∆^5^-desaturase (*82*)  Substrate: lathosterol  ∆^24^ -Sterolreductase (*82*)  Substrate: cholesta-5,7,24-trienol | – | Cholesterol precursor, VD3 precursor (*82*). |
| 2 | 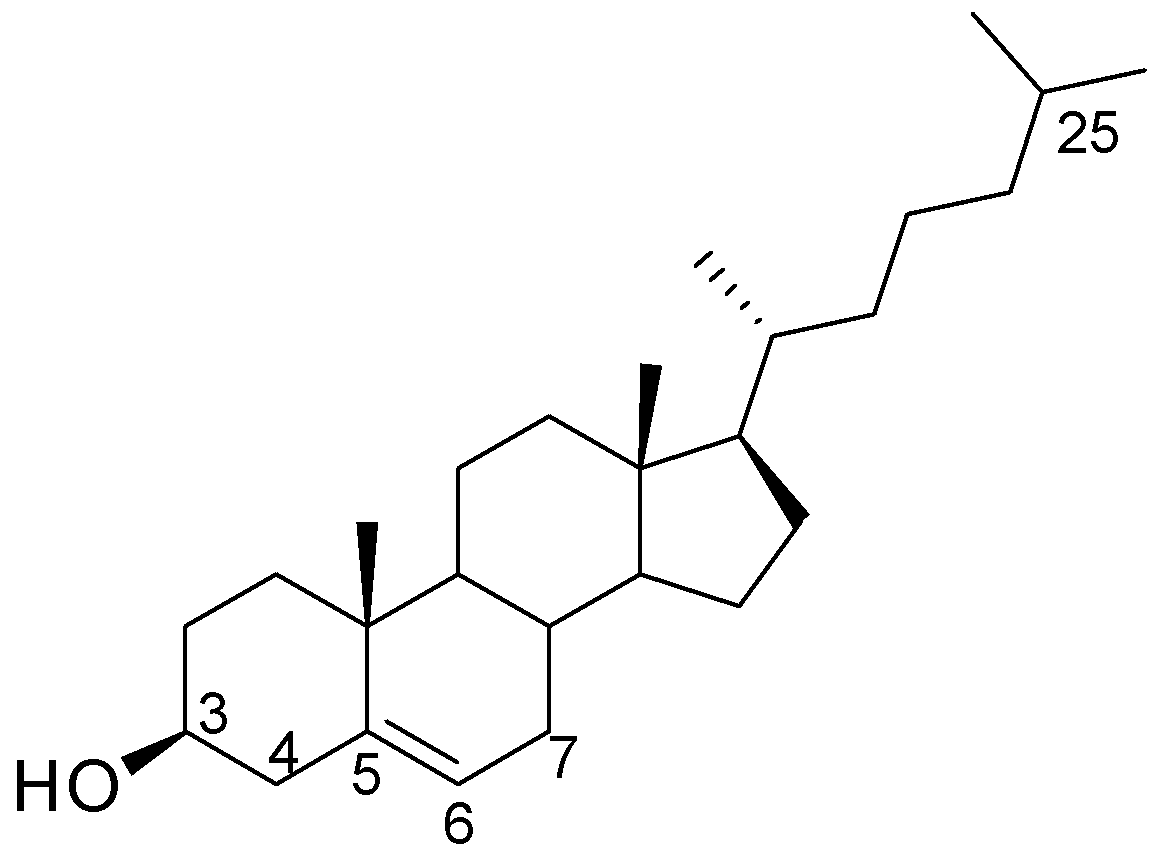  cholesterol | 7-dehydrocholesterol reductase (DHCR7) (*82*)  Substrate: 7DHC  ∆^24^ –Sterol reductase (*82*)  Substrate: desmosterol | ERRα (*83*) | Precursor of steroid hormones and bile acids (*84*). |
| 3 | 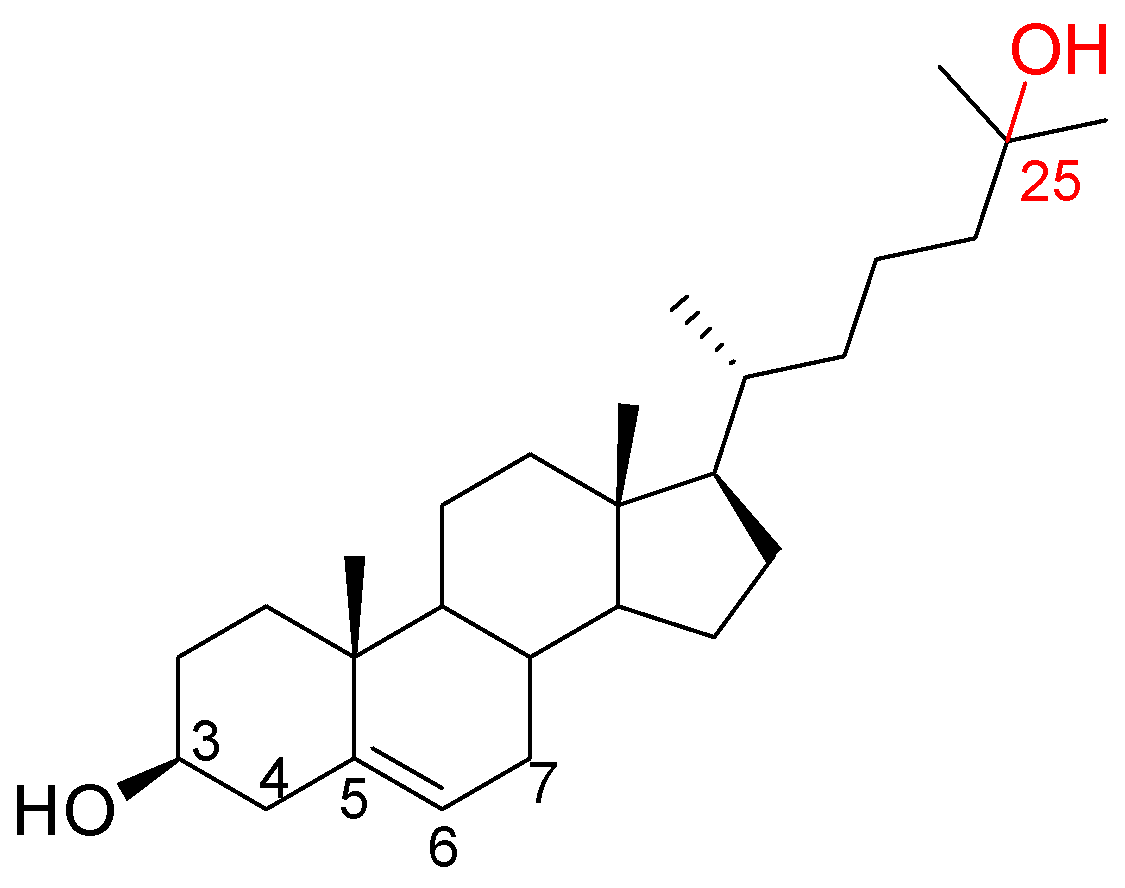  25-hydroxy-cholesterol  (25OHC) | Cholesterol-25-hydroxylase (CH25H) (*13*)  Substrate: cholesterol | LXRs (*85*) | Regulation of cholesterol biosynthesis, pro- and anti-inflammatory activity (*13*). |
| 4 | 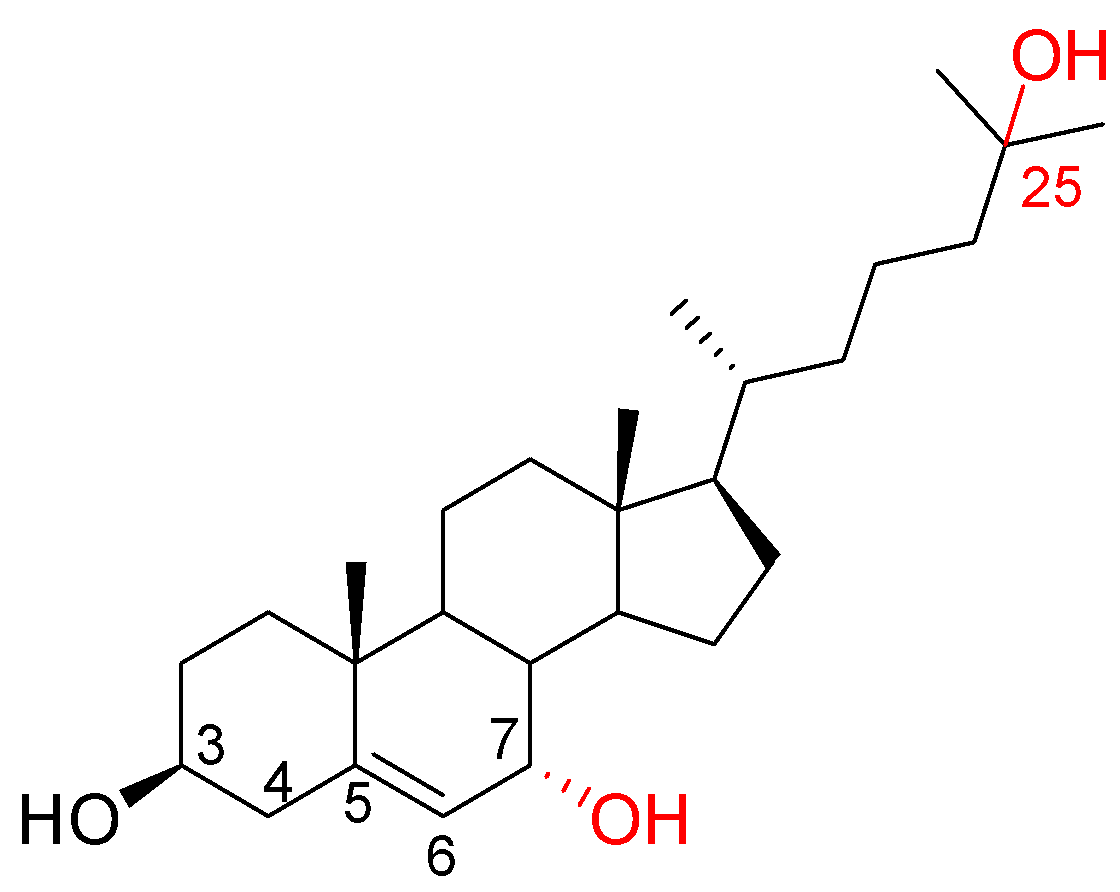  7α,25-dihydroxy-cholesterol  (7α,25diOHC) | CYP7B1 (*49*)  Substrate:  25OHC | EBI2 (*86*) | Lymphocyte migration (*86*). |
| 5 | 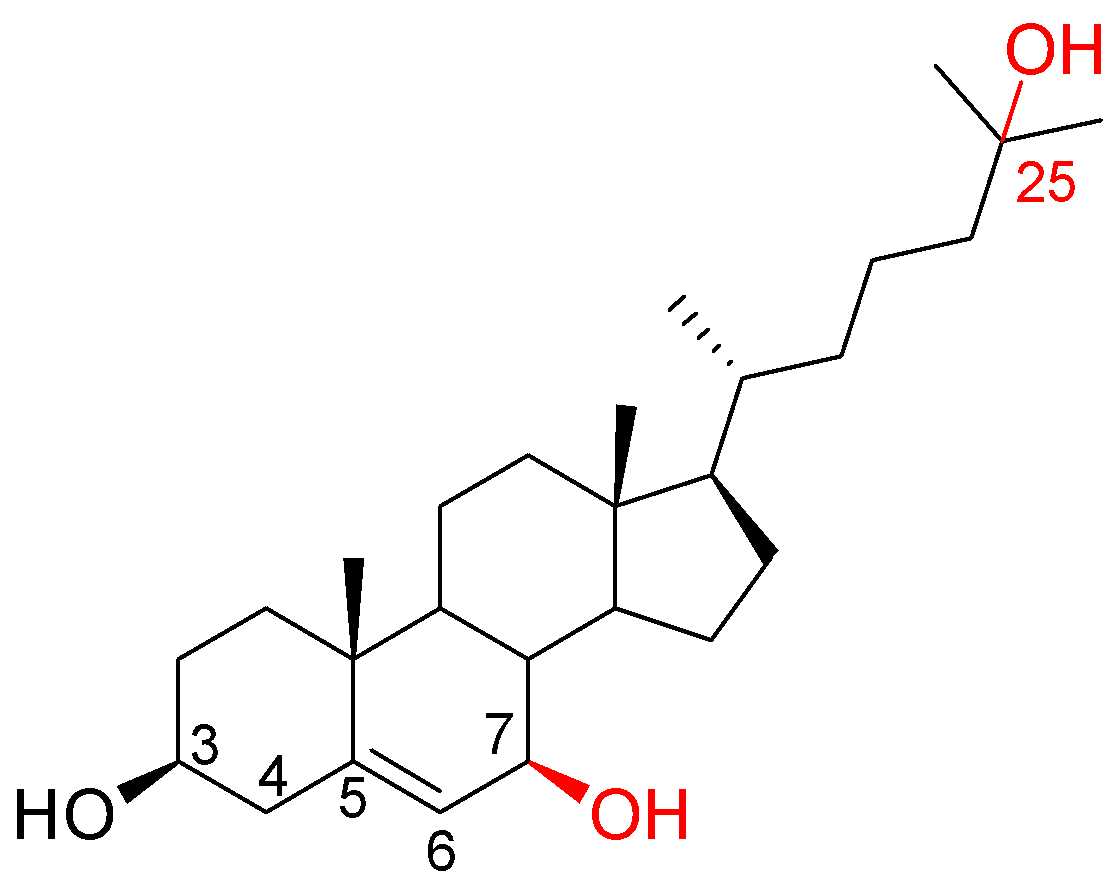  7β,25-dihydroxy-cholesterol  (7β,25diOHC) | ROS (*87*)  Substrate:  25OHC | EBI2 (*17*) | Lymphocyte migration (*17*). |
| 6 | 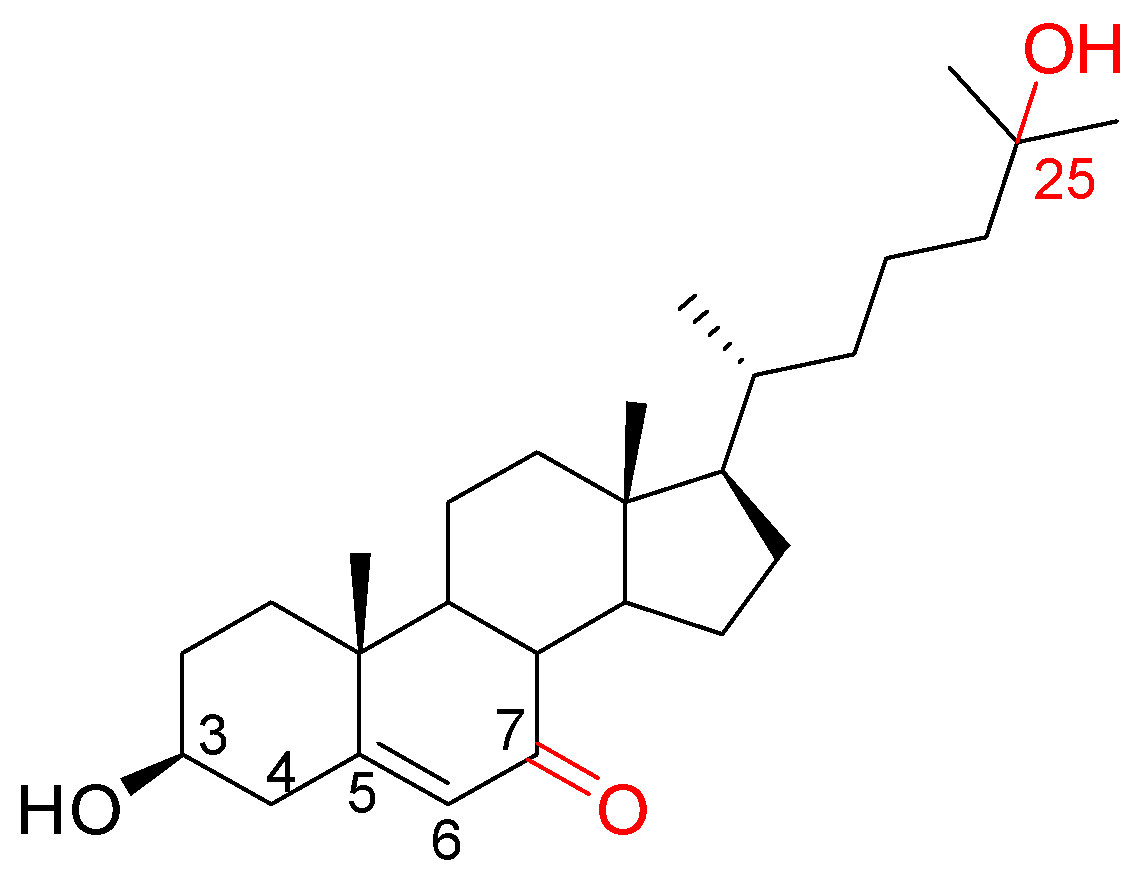  7keto-25-hydroxy-cholesterol  (7keto,25OHC) | Hypothetical: 7α-hydroxysteroid dehydrogenase (from gut bacteria, 7αHSD) (*32*)  Substrate: 7α,25diOHC, 7β,25diOHC  Hypothetical: ROS (*87*)  Substrate: 25OHC | – | – |
| 7 | 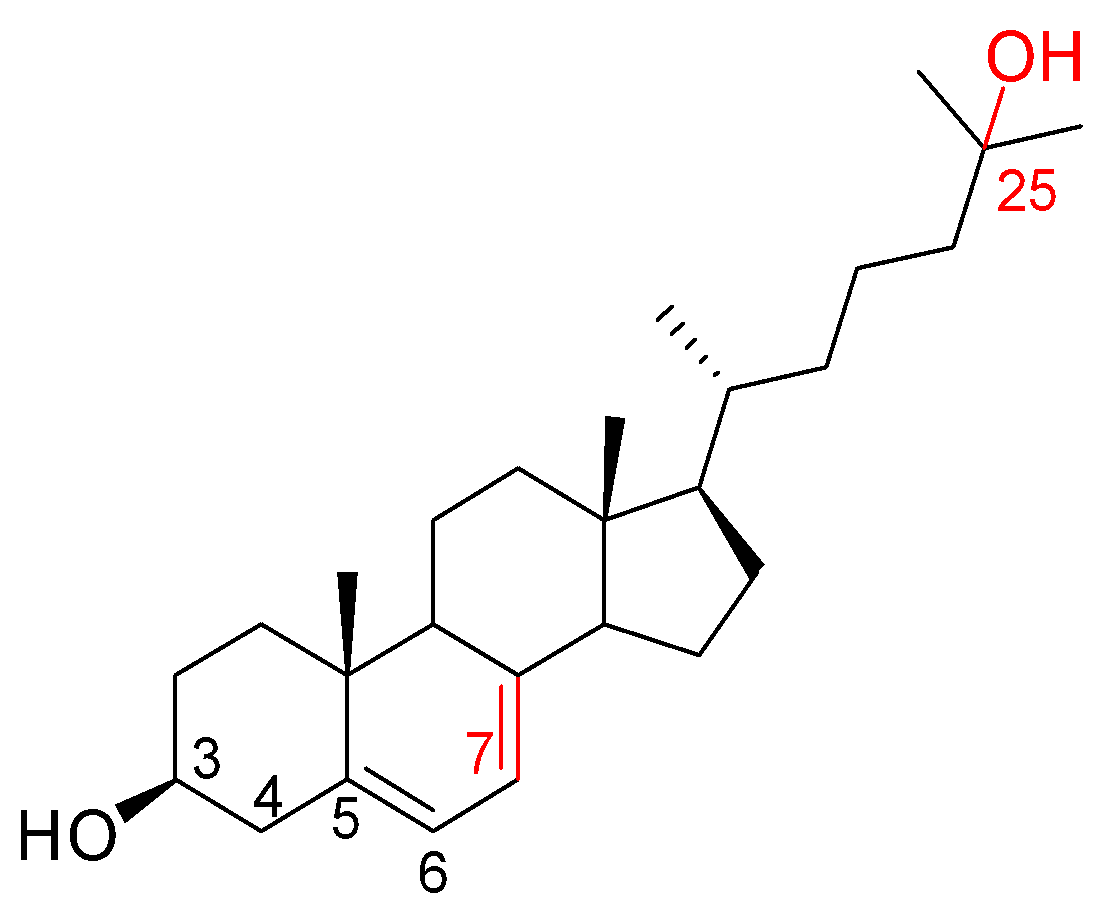  25-hydroxy-7-dehydrocholesterol  (25OH7DHC) | Hypothetical:  CH25H  Substrate: 7DHC | – | – |
| 8 | 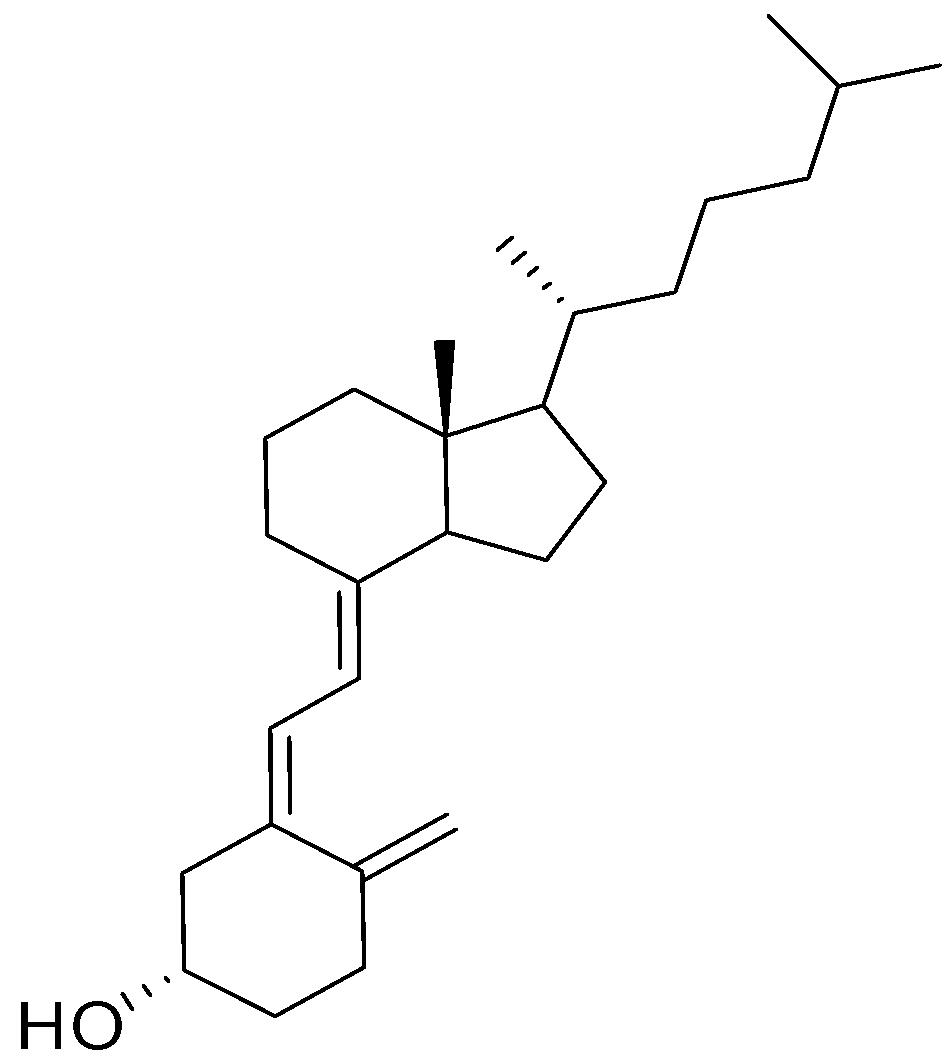  vitamin D_3_  (VD3) | UV-B exposure (*88*)  Substrate: 7DHC | – | Physiologically inactive precursor of active forms of VD3 (*89*). |
| 9 | 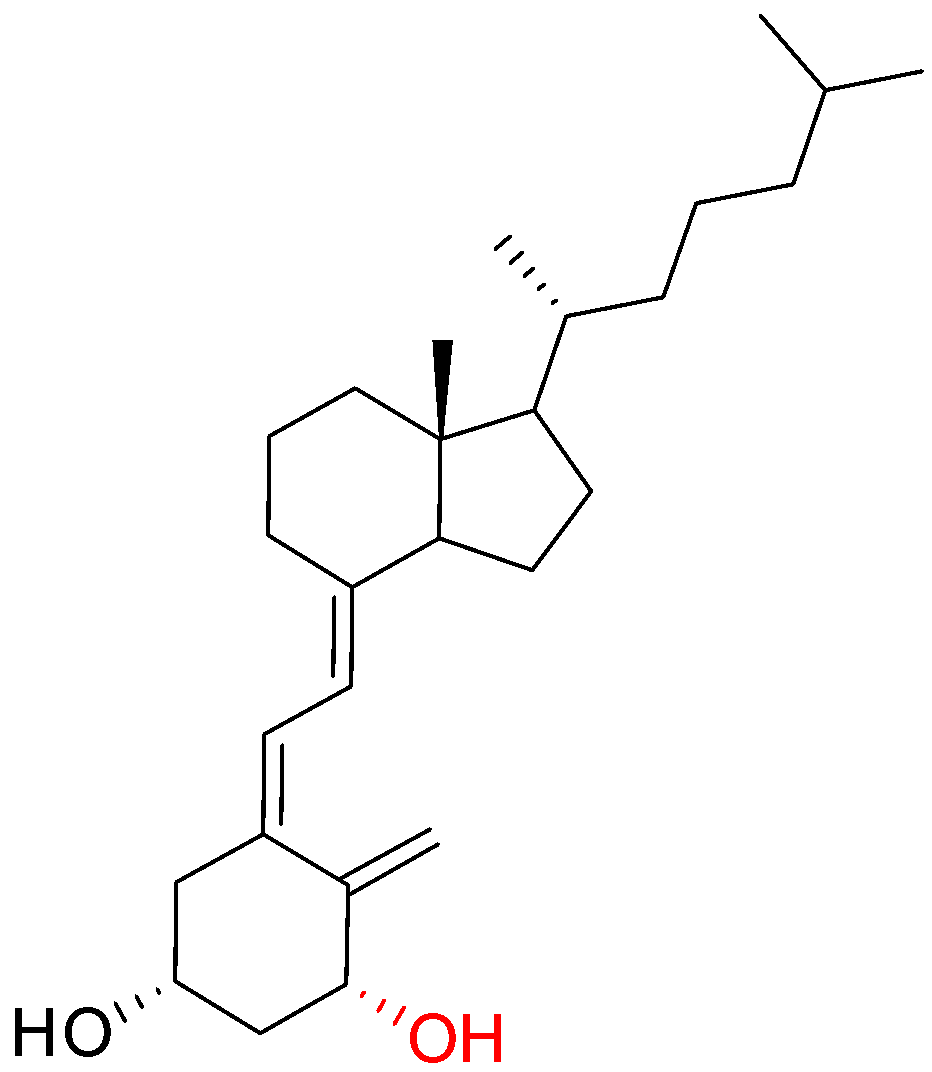  1α-hydroxy-VD3  (1αOHVD3, alfacalcidol) | Synthetic secosteroid (*90*) | VDR (alfa-calcidol is VD3 mimetic) (*91*) | Modulate expression of major monocyte antigens CD14 and HLA-DR, and receptor TLR2, involved in antigen recognition and processing (*90*). |
| 10 | 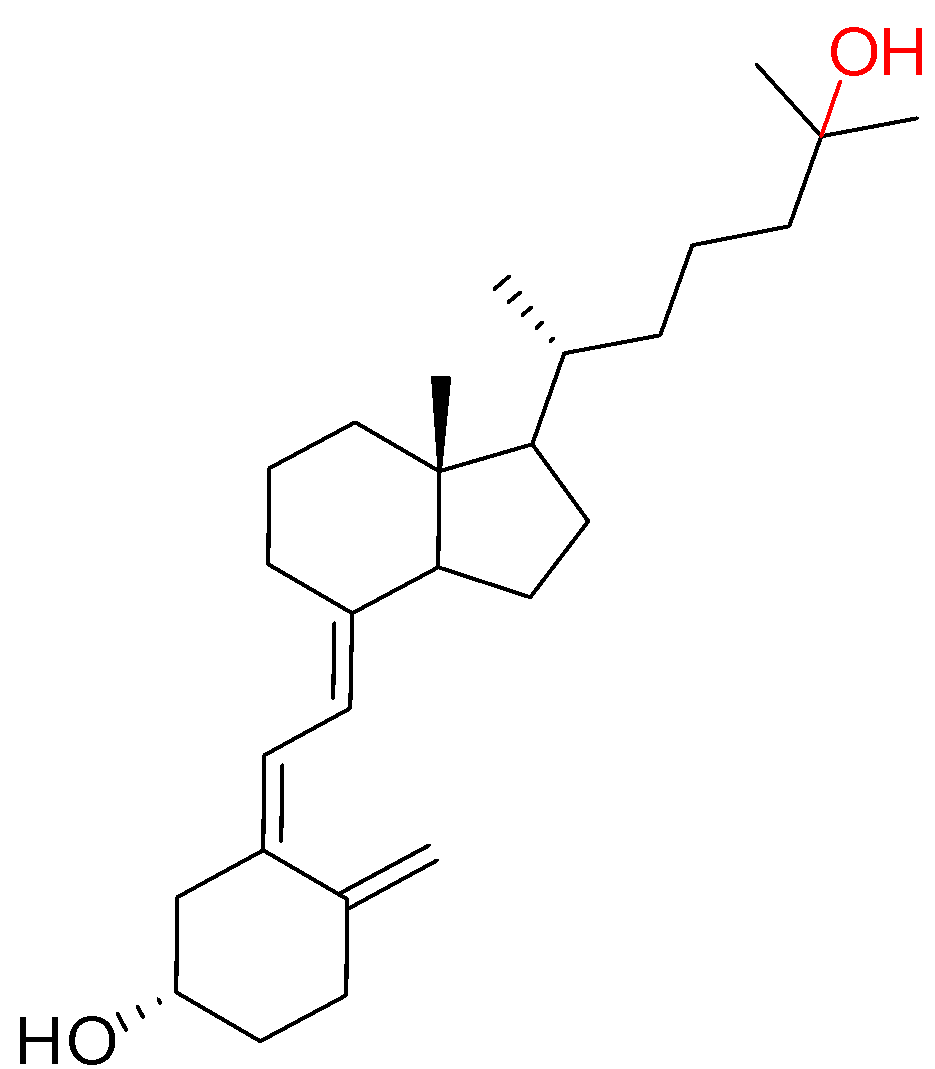  25-hydroxy-VD3  (25OHVD3) | CYP2R1 (*92*)  Substrate:  VD3 |  | Intermediate of VD3 biosynthesis (*93*). |
| 11 | 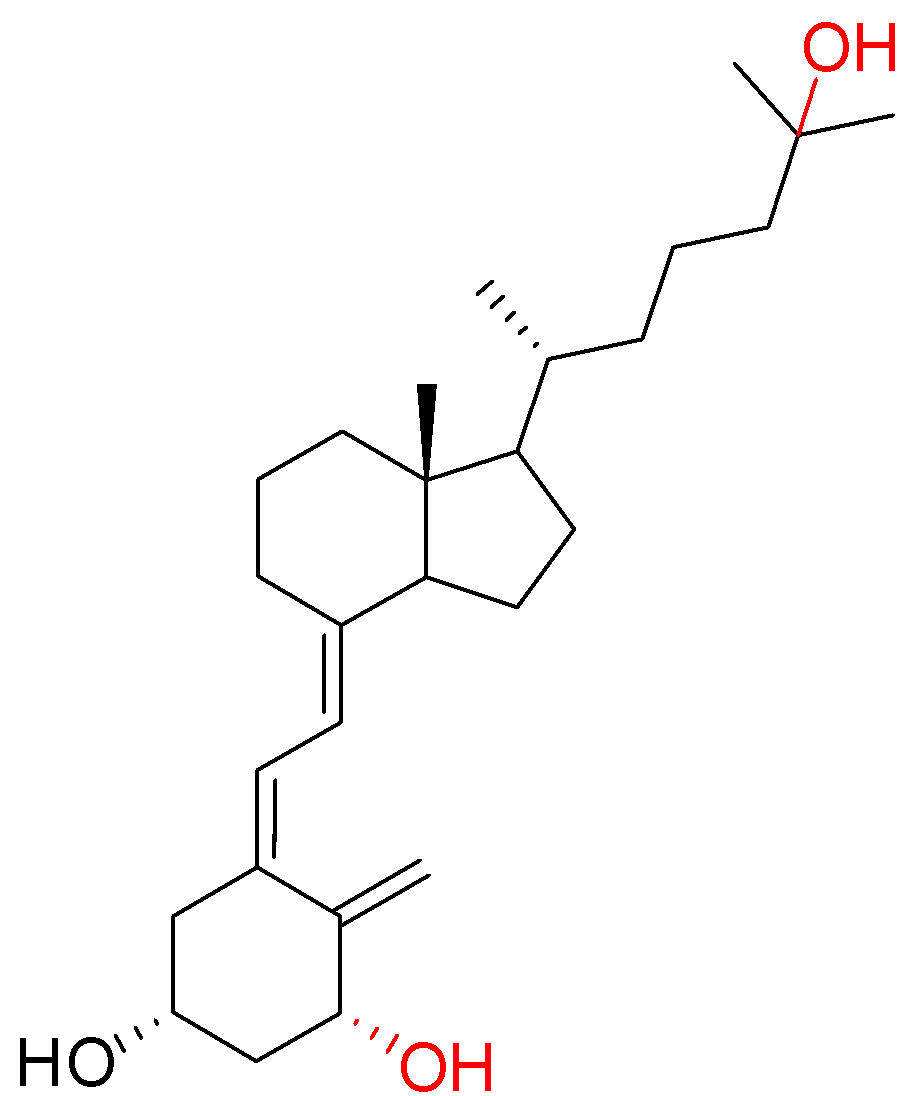  1α,25-dihydroxy-VD3  (1α,25diOHVD3) | CYP27B1 (*94*)  Substrate:  25OHVD3  CYP2R1 (*92*)  Substrate:  1αOHVD3 | VDR (*89*) | Induces expression of antimicrobial factors (*95*).  Stimulates differentiation of monocytic precursors into macrophage-like cells (*94*). Has immunosuppressive activity via dendritic cell differentiation and maturation inhibition (*96*). |
| 12 | 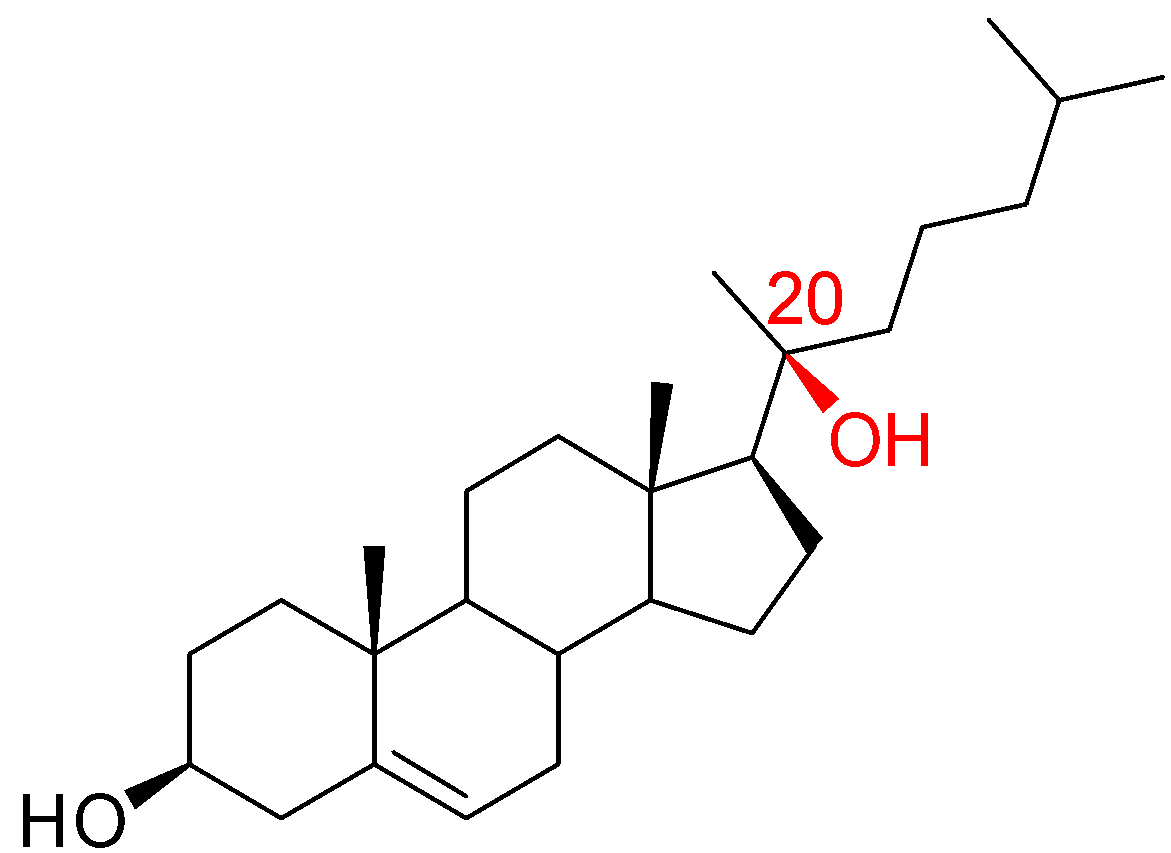  20S-hydroxy-cholesterol  (20SOHC) | CYP11A1 (*97*)  Substrate: cholesterol | SMO (*98*) | Activator of Sonic Hedgehog (*98*) and Notch signaling pathways. Stimulates osteogenic differentiation of pluripotent mesenchymal stem cells and inhibits their adipogenic differentiation (*99*). |
| 13 | 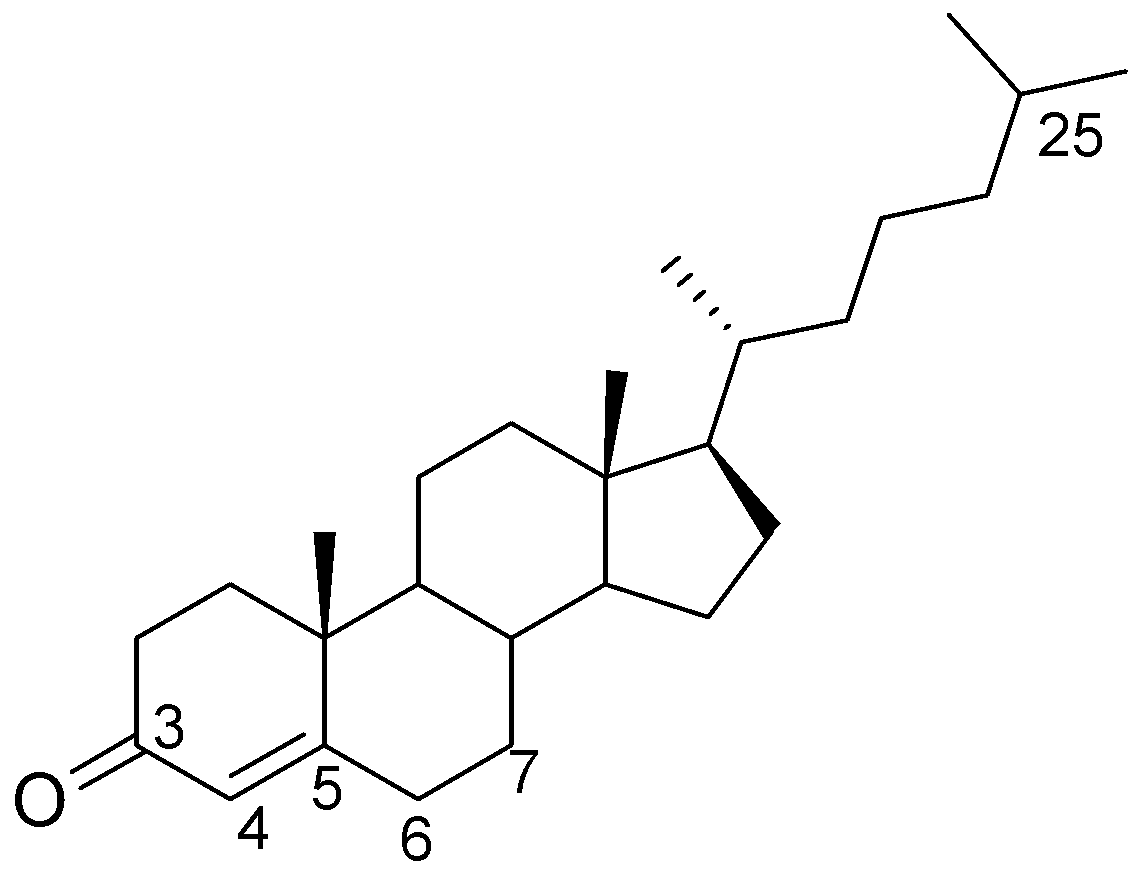  cholest-4-en-3-one  (cholestenone) | Cholesterol oxidase (from gut bacteria) (*100*)  3β-hydroxysteroid dehydrogenase (3βHSD)  Substrate: cholesterol | - | Intermediate of biosynthesis of steroids (*93*). |
| 14 | 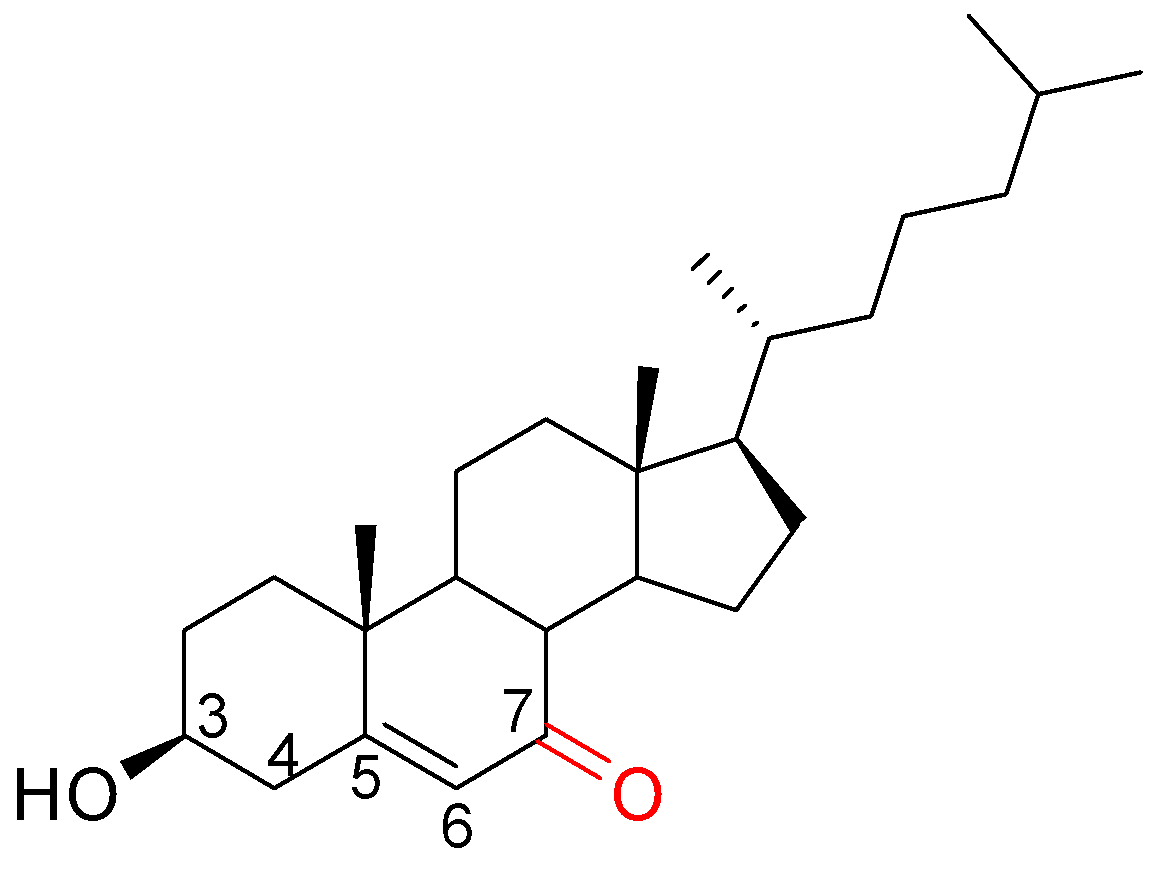  7-ketocholesterol  (7ketoC) | CYP7A1 (*101*)  Substrate: 7DHC  ROS (*87*)  Substrate: cholesterol | TLR4 (*102*) | Involved in bile acid biosynthesis (110). Inhibit HMG-CoA reductase, thus modulating cholesterol biosynthesis (*103*). Inhibits CYP7A1 (*104*). Induces inflammation through TLR4 (*102*). Induces apoptosis (*105*). |

#### Table S2. Binding affinity parameters for various ligands to Mtb CYP124, CYP125, and CYP142 based on spectrophotometric titration assay

| **Ligand** | **Kd_app_ ± SE (μM)** | | |
| --- | --- | --- | --- |
|  | **CYP124** | **CYP125** | **CYP142** |
| Cholestenone | 3.4 ± 0.6 | 10.8 ± 1.8 | 2.8 ± 0.4 |
| 25OHC-one | ND | ND | NS |
| 20SOHC | 4.6 ± 0.7 | ND | ND |
| 7βOHC-one | 13.2 ± 1.7 | 26 ± 3 | 27 ± 8 |
| 5-cholesten-3β,25-diol-7-one | ND | ND | NS |
| 25OHC | ND | NS | ND |
| 7βOHC | 11.1 ± 1.3 | 13 ± 2 | 6.0 ± 1.4 |
| 7DHC | 4.9 ± 1.5 | ND | ND |
| 7ketoC | 6.5 ± 0.8 | 12.4 ± 1.7 | 4.3 ± 0.8 |
| 7α,25diOHC | ND | ND | ND |
| 25OH7DHC | ND | ND | ND |
| VD3 | 18 ± 2 | NS | ND |
| VD2 | ND | NS | NS |
| 1αOHVD3 | 34 ± 4 | ND | ND |
| 1αOHVD2 | 14 ± 2 | ND | NS |
| 25OHVD3 | ND | 3.8 ± 0.6 | ND |
| 1α,25diOHVD3 | ND | NS | ND |
| CHpImi | 3.1 ± 0.5 | 1.61 ± 0.03 | 31 ± 3 |
| CHImi | 1.45 ± 0.08 | 31 ± 4 | 3.73 ± 0.19 |
| Cpd5' | 1.93 ± 0.16 | 16 ± 5 | 0.17 ± 0.06 |

Kd_app_ values of ligand-binding to CYPs were determined by spectrophotometric titration using a dual-beam spectrophotometer with 1 or 2 μM of the CYP (see Fig. S1, S2, and S3). The data represent the mean ± s.e. for at least three independent experiments. NS – no spectral shift in the Soret band was observed. ND – Kd_app_ value could not be determined due to lack of spectral changes.

#### Table S3. Crystallographic data collection and refinement statistics

| **Data collection** | Cholestenone | VD3 | 1αOHVD3 | CHImi | Cpd5' |
| --- | --- | --- | --- | --- | --- |
| PDB ID code | 6T0F | 6T0G | 6T0H | 6T0K | 6T0L |
| Space group | P12_1_1 | P12_1_1 | P12_1_1 | P12_1_1 | P4_3_2_1_2 |
| **Cell dimensions** |  |  |  |  |  |
| a, b, c (Å) | 93.67, 81.27, 155.85 | 51.58, 74.81, 56.59 | 51.56, 74.97, 56.63 | 51.62, 75.03, 56.52 | 71.57, 71.57, 195.61 |
| α, β, γ (°) | 90, 107.286, 90 | 90, 107.058, 90 | 90, 106.952, 90 | 90, 106.924, 90 | 90, 90, 90 |
| Wavelength (Å) | 1.03320 | 0.98400 | 0.97625 | 0.97800 | 0.96600 |
| No. of observations | 3655547 (273962) | 498997 (30487) | 494466 (28827) | 550957 (27897) | 706499 (24462) |
| No. of unique reflections | 360719 (26345) | 164106 (11943) | 161384 (11371) | 183003 (12593) | 60287 (3973) |
| Resolution (Å) | 50 - 1.49  (1.53 - 1.49) | 30 - 1.10  (1.13 - 1.10) | 20 - 1.10  (1.13 - 1.10) | 30 - 1.06  (1.09 - 1.06) | 30 - 1.66  (1.70 - 1.66) |
| Rmeas | 0.172 (4.341) | 0.195 (2.013) | 0.135 (2.889) | 0.110 (2.348) | 0.120 (3.557) |
| Rpim | 0.054 (1.332) | 0.108 (1.166) | 0.074 (1.730) | 0.060 (1.409) | 0.034 (1.395) |
| I/σI | 8.21 (0.36) | 3.65 (0.36) | 4.49 (0.35) | 5.64 (0.35) | 12.05 (0.47) |
| CC1/2 | 99.9 (13.6) | 99.0 (13.3) | 99.1 (10.1) | 99.6 (12.3) | 99.9 (40.1) |
| Completeness (%) | 99.2 (98.3) | 98.8 (97.5) | 96.8 (92.7) | 98.3 (91.2) | 98.7 (89.4) |
| Redundancy | 10.1 (10.4) | 3.0 (2.6) | 3.1 (2.5) | 3.0 (2.2) | 11.7 (6.2) |
| **Refinement statistics** |  |  |  |  |  |
| Resolution (Å) | 50 - 1.65  (1.69 - 1.65) | 20 - 1.30  (1.33 - 1.30) | 20 - 1.18  (1.21 - 1.18) | 30 - 1.18  (1.21 - 1.18) | 30 - 1.80  (1.85 - 1.80) |
| Number of reflections (total/unique) | 2704584/266740 | 318171/99588 | 414682/131383 | 424610/133380 | 706499/60287 |
| Rwork/Rfree | 0.1558/0.1835 | 0.1631/0.1903 | 0.1559/0.1811 | 0.1308/0.1565 | 0.1633/0.1923 |
| CC* in highest shell | 0.829 (18885) | 0.817 (6321) | 0.759 (9345) | 0.813 (8770) | 0.943 (3376) |
| CCwork/CCfree in highest shell | 0.789/0.744 | 0.760/0.687 | 0.637/0.589 | 0.744/0.646 | 0.894/0.867 |
| Number of atoms |  |  |  |  |  |
| Protein | 15010 | 3695 | 4114 | 4119 | 3531 |
| Heme | 215 | 43 | 43 | 43 | 43 |
| Ligand | 112 | 20 | 49 | 32 | 16 |
| Solvent | 2734 | 810 | 812 | 724 | 404 |
| Number of TLS groups | 12 | - | - | - | 2 |
| B-factors (Å^2^) |  |  |  |  |  |
| Protein | 26.2 | 13.7 | 15.4 | 14.6 | 37.7 |
| Heme | 18.7 | 8.8 | 10.8 | 9.5 | 24.9 |
| Ligand | 19.1 | 15.9 | 19.1 | 13.3 | 33.5 |
| Solvent | 39.5 | 28.5 | 31.6 | 33.2 | 45.7 |
| R.m.s.d |  |  |  |  |  |
| Bond lengths (Å) | 0.012 | 0.012 | 0.008 | 0.007 | 0.011 |
| Bond angles (°) | 1.17 | 1.18 | 1.08 | 0.89 | 1.05 |
| Ramachandran statistics |  |  |  |  |  |
| Favoured (%) | 97.61 | 97.88 | 98.13 | 97.67 | 98.57 |
| Allowed (%) | 2.39 | 2.12 | 1.87 | 2.33 | 1.43 |

#### Table S4. Transcriptional changes of proteins potentially involved in host steroid transport and metabolism triggered by overexpression of TFs

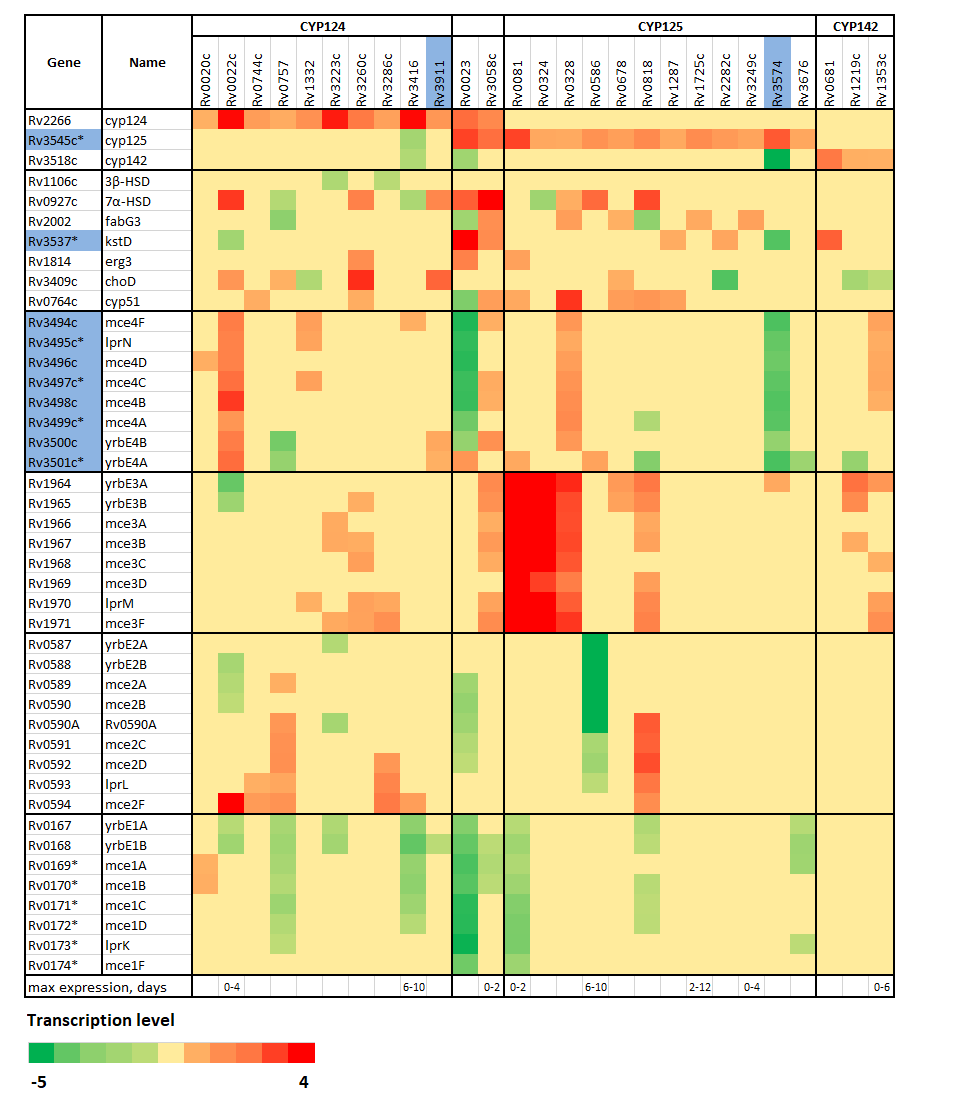

This table is based on (*27*) and includes steroid-metabolizing CYPs CYP124, CYP125, and CYP142, steroid-metabolizing enzymes 3βHSD, 7αHSD, fabG3, Rv3537, erg3, choD, and CYP51, and genes of mce loci. Genes essential for *in vitro* growth on cholesterol are marked with blue cells (*10*), and genes essential for *in vivo* growth are marked with asterisks (*****) (*106*). Co-expression of CYP124 and mce4 genes is observed, as well as co-expression of CYP125 and mce3 genes. Transporter loci mce4, mce3, and mce2 are co-activated with CYPs and steroid-metabolizing enzymes, while transporter locus mce1 is mainly repressed under overexpression of the TFs. 3βHSD (Rv1106c) catalyzes the first step in the cholesterol metabolism pathway, converting cholesterol to the 4‐en‐3‐one (*107*); 7αHSD catalyzes the dehydrogenation of a hydroxyl group at position 7 of the steroid skeleton of bile acids in *Escherichia coli* (*32*); fabG3 (possible 3α, 20β-hydroxysteroid dehydrogenase FabG3/cortisone reductase/(R)-20-hydroxysteroid dehydrogenase) probably plays an role in steroid metabolism in Mtb (*33*); *kstD (Rv3537)* catalyzes the elimination of the C-1 and C-2 hydrogen atoms of the A-ring from the polycyclic ring structure of 3-ketosteroids, also involved in the formation of 3-keto-1,4-diene-steroid from 3-keto-4-en-steroid (*34*); erg3 (Rv1814) – membrane-bound C-5 sterol desaturase, that in *Candida albicans* has been shown to confer azole resistance (*108*); choD (Rv3409c) – cholesterol oxidase catalyzing the same reaction as 3βHSD, which is important for virulence and *in vivo* growth, but not essential for cholesterol degradation (*10*, *109*); cyp51 is a lanosterol 14-α demethylase with unknown natural substrate (*110*). The mce4 operon imports cholesterol. The natural substrates of the mce2 and mce3 transporters have not yet been identified. The mce1 operon may serve as a mycolic acid re-importer (*111*). Rv0020c (fhaA) regulates cell growth and peptidoglycan synthesis by binding to MviN and may inhibit the late stages of peptidoglycan synthesis (*112*); Rv0023 is a possible transcriptional regulatory protein that is up-regulated 12-fold in 7 days after infection of human macrophages (*113*); Rv0081 is a member of the dormancy regulon induced in response to hypoxia, low levels of nitric oxide, and carbon monoxide, and serving as an immunogenic antigen (*114*); Rv3223c (sigH) is an alternative RNA polymerase sigma-E factor that plays a role in the oxidative-stress response (*115*); Rv3416 (whiB3) is a redox-sensing transcriptional regulator that maintains intracellular redox homeostasis by regulating metabolism of pathogenic lipids and of polyketide biosynthesis (*116*).

#### Table S5. Activity of cpd5' against Mtb H37Rv residing in macrophages

| [cpd5'], μg/ml | 3[H]-uracil uptake  CPM ± SD × 10^3^ | Reduction of mycobacterial growth, % |
| --- | --- | --- |
| 0 | 76 ± 5 | 0 |
| 0.63 | 75 ± 3 | 2 |
| 2.5 | 57 ± 6 | 25 |
| 10.0 | 20 ± 3 | 76 |
| 40.0 | 15.2 ± 1.6 | 82 |

**Supplementary Methods**

**Synthesis of 3β,25-dihydroxycholesta-5,7-diene (25OH7DHC).**

The scheme of the synthesis is shown in Fig. S8. Melting points were measured using a Boetius apparatus and are not corrected. UV spectra were recorded in MeOH on a Specord UV/Vis instrument. IR spectra were obtained from KBr pellets or films using a UR-20 instrument. ^1^H NMR and ^13^C NMR spectra were recorded in CDCl_3_ or C_5_D_5_N solutions using residual solvent peaks as internal standards (δ_Н_ 7.26 ppm and δ_С_ 77.16 ppm for CDCl_3_; δ_Н_ 7.58 ppm and δ_С_ 135.9 ppm for С_5_D_5_N) on a Bruker AVANCE-500 instrument (operating frequency 500 MHz for ^1^H and 125 MHz for ^13^C). Positive-ion mass spectra were obtained on an LCQ Fleet mass spectrometer (Thermo Electron Corp.) using electrospray ionization (ESI). Ozone was generated from an oxygen cylinder in an ozonator with output of 2 grams of ozone per hour. The course of reactions was monitored by TLC on Kieselgel 60 F254 plates with detection by anisaldehyde followed by heating or under a UV lamp. Reaction mixtures were separated by chromatography on silica gel 40/60 (Kieselgel 60, Merck).

*3β-Acetoxyergosta-5,7,22-trien (***2***).* Ergosterol **1 (**1 g, 2.53 mmol) was dissolved in dry pyridine (4 ml). To this solution, acetic anhydride (2 ml, 21 mmol) and 4-dimethylaminopyridine (30 mg, 0.25 mmol) were added. The mixture was stirred at room temperature for 10 h. The reaction mixture was diluted with water (40 ml), and the precipitated crystals were separated by filtration, washed with water, and purified by recrystallization from ethanol to give yellowish crystals of 3-acetate **(2)** (791 mg, 70%); mp 150-152 °С. IR (KBr) ν, cm^−1^: 2975, 1725, 1215, 1045. ^1^H NMR (CDCl_3_, 500 MHz): δ_H_ 5.56 (m, 1H), 5.38 (m, 1H), 5.14-5.25 (m, 2H), 4.71 (m, 1H), 2.50 (m, 1H), 2.36 (m, 1H), 2.04 (s, 3H), 1.03 (d, *J* = 6.5 Hz, 3H), 0.95 (s, 3H), 0.91 (d, *J* = 6.8 Hz, 3H), 0.80-0.86 (m, 6H), 0.62 (s, 3H). ^13^C NMR (CDCl_3_, 125 MHz): δ_C_ 170.7 (s), 141.7 (s), 138.7 (s), 135.7 (d), 132.1 (d), 120.3 (d), 116.4 (d), 72.9 (d), 55.8 (d), 54.6 (d), 46.1 (d), 42.9 (s, d), 40.5 (d), 39.1 (t), 38.0 (t), 37.2 (s), 36.8 (t), 33.2 (d), 28.4 (t), 28.2 (t), 23.1 (t), 21.5 (q), 21.2 (q), 21.1 (t), 20.1 (q), 19.8 (q), 17.7 (q), 16.3 (q), 12.2 (q). MS (ESI^+^), *m/z* (*I_rel_*, %): 379 ([M-AcOH+H]^+^, 100), 301 (45), 163 (62).

*3β-Acetoxy-5α,8α-(1,4-dioxo-1,2,3,4-tetrahydrophthalazine-2,3-diyl)ergosta-6,22-diene (***3***).* To a solution of diene **2** (595 mg, 1.36 mmol) and phthalic hydrazide (740 mg, 4.57 mmol) in dry CH_2_Cl_2_ (20 ml), a solution of Pb(OAc)_4_ (2 g, 4.57 mmol) and CH_3_COOH (100 μl) in dry CH_2_Cl_2_ (10 ml) was added dropwise at 0 °C for 1 h under argon. The resulting mixture was stirred at room temperature for 1 h, and the precipitate was removed by filtration. The filtrate was washed with water and dried over Na_2_SO_4_, concentrated *in vacuo*, and purified by column chromatography on silica gel (petroleum ether/EtOAc = 15:1=> 5:1) to give Diels-Alder adduct **3** (760 mg, 94%) as a yellow amorphous solid; mp 108-112°С. IR (KBr) ν, cm^−1^: 2960, 2870, 1745, 1725, 1310, 1245. ^1^H NMR (CDCl_3_, 500 MHz): δ_H_ 8.12 (m, 2H), 7.68 (m, 2H), 6.65 (d, *J* = 8.3 Hz, 1H), 6.28 (d, *J* = 8.3 Hz, 1H), 5.18 (m, 2H), 4.70 (m, 1H), 4.00 (dd, *J* = 13.9, 3.8 Hz, 1H), 3.93 (dd, *J* = 13.9, 3.8 Hz, 1H), 2.01 (s, 3H), 1.04 (s, 3H), 1.01 (d, *J* = 6.6 Hz, 3H), 0.89 (d, *J* = 6.9 Hz, 3H), 0.83 (d, *J* = 6.8 Hz, 3H), 0.82 (s, 3H), 0.81 (d, *J* = 6.8 Hz, 3H). ^13^C NMR (CDCl_3_, 125 MHz): δ_C_ 170.1 (s), 162.1 (s), 159.7 (s), 137.7 (d), 135.2 (d), 132.8 (d), 132.8 (d), 132.1 (d), 130.5 (s), 129.9 (s), 129.1 (d), 127.0 (d), 126.9 (d), 69.9 (d), 68.2 (s), 66.8 (s), 56.5 (d), 50.3 (d), 48.8 (d), 44.1 (s), 42.7 (d), 40.4 (s), 39.9 (d), 39.2 (t), 34.9 (t), 33.0 (d), 31.1 (t), 28.1 (t), 26.0 (t), 24.4 (t), 21.7 (t), 21.3 (q), 20.8 (q), 19.9 (q), 19.6 (q), 18.3 (q), 17.4 (q), 13.3 (q). MS (ESI^+^), *m/z* (*I_rel_*, %): 639 [M+K] ^+^ (5), 621 [M+Na] ^+^ (100), 599 [M+H] ^+^ (52).

*3β-Acetoxy-5α,8α-(1,4-dioxo-1,2,3,4-tetrahydrophthalazine-2,3-diyl)-22-hydroxy-23,24-bisnorchol-6-ene (****4****).* A mixture of O_3_ and O_2_ was bubbled through a solution of Diels-Alder adduct **3** (760 mg, 1.27 mmol) in dry CH_2_Cl_2_ (40 ml) and dry pyridine (0.2 ml) at –60 °C for 1 h. After almost quantitative conversion of the starting material, stirring was continued for 5 min followed by the addition of NaBH_4_ (241 mg, 6.35 mmol). Then the mixture was stirred at room temperature for 3 h. The reaction mixture was washed with 0.5 M HCl (30 ml) and brine (60 ml). The organic phase was dried over Na_2_SO_4_, the solvent was evaporated, and the residue was purified by column chromatography on silica gel (petroleum ether/EtOAc = 10:1) to give alcohol **4** (401 mg, 63%) as a colorless amorphous solid; mp 143-145 °С. IR (KBr) ν, cm^−1^: 3455, 2960, 2855, 1740, 1725, 1315, 1250. ^1^H NMR (CDCl_3_, 500 MHz): δ 8.12 (m, 2H), 7.69 (m, 2H), 6.66 (d, *J* = 8.2 Hz, 1H), 6.28 (d, *J* = 8.2 Hz, 1H), 4.69 (m, 1H), 3.97 (m, 2H), 3.64 (dd, *J* =10.1, 7.3 Hz, 1H), 3.37 (dd, *J* =10.1, 7.3 Hz, 1H), 2.01 (s, 3H), 1.05 (d, *J* = 7.4 Hz, 3H), 1.04 (s, 3H), 0.85 (s, 3H). ^13^C NMR (CDCl_3_, 125 MHz): δ 170.1 (s), 162.1 (s), 159.8 (s), 137.9 (d), 132.9 (d), 132.8 (d), 130.4 (s), 129.9 (s), 128.9 (d), 127.0 (d), 126.9 (d), 69.9 (d), 68.2 (s), 67.6 (t), 66.8 (s), 53.3 (d), 50.3 (d), 48.5 (d), 44.4 (s), 40.4 (s), 39.1 (t), 38.3 (d), 34.9 (t), 31.1 (t), 27.2 (t), 26.0 (t), 24.4 (t), 21.8 (t), 21.3 (q), 18.3 (q), 16.6 (q), 13.2 (q). MS (ESI^+^), *m/z* (*I_rel_*, %): 1087 [2M+Na] ^+^ (9), 533 [M+H] ^+^ (100).

*3β-Acetoxy-5α,8α-(1,4-dioxo-1,2,3,4-tetrahydrophthalazine-2,3-diyl)-22-iodo-23,24-bisnorchol-6-ene (***5***).* Iodine (381 mg, 1.50 mmol) was added to a stirred, cooled to 0 °C solution of imidazole (204 mg, 3.00 mmol) and triphenylphosphine (393 mg, 1.50 mmol) in dry CH_2_Cl_2_ (20 ml). The mixture was stirred at 0 °C for 15 min and treated with a solution of alcohol **4** (401 mg, 0.75 mmol) in dry CH_2_Cl_2_ (5 ml) for 10 min. Stirring was continued at 5 °C for 0.5 h and then at room temperature for 2.0 h. The precipitate was removed by filtration, and washed with CH_2_Cl_2_ (12 ml), and then the combined filtrates were washed subsequently with 2% Na_2_S_2_O_3_ (30 ml), 0.1 M HCl (10 ml), and brine (20 ml), and then dried over Na_2_SO_4_. The solvent was evaporated and the residue was purified by column chromatography on silica gel (petroleum ether/EtOAc = 19:1) to give 22-iodide **5** (450 mg, 93%) as a yellow amorphous solid; mp 138-140 °С. IR (KBr) ν, cm^−1^: 2950, 1750, 1725, 1310, 1245. ^1^H NMR (CDCl_3_, 500 MHz): δ_H_ 8.13 (m, 2H), 7.69 (m, 2H), 6.66 (d, *J* = 7.9 Hz, 1H), 6.27 (d, *J* = 7.9 Hz, 1H), 4.69 (m, 1H), 3.98 (m, 2H), 3.34 (dd, *J* = 10.1, 2.6 Hz, 1H), 3.11 (dd, *J* = 10.1, 2.6 Hz, 1H), 2.01 (s, 3H), 1.05 (d, *J* = 6.8 Hz, 3H), 1.04 (s, 3H), 0.86 (s, 3H). ^13^C NMR (CDCl_3_, 125 MHz): δ_C_ 170.1 (s), 162.1 (s), 159.7 (s), 137.9 (d), 132.9 (d), 132.8 (d), 130.3 (s), 129.8 (s), 128.8 (d), 127.0 (d), 126.8 (d), 69.8 (d), 68.1 (s), 66.7 (s), 55.7 (d), 50.2 (d), 48.5 (d), 44.3 (s), 40.3 (s), 38.9 (t), 36.7 (d), 34.8 (t), 31.0 (t), 27.1 (t), 26.0 (t), 24.3 (t), 21.6 (t), 21.3 (q), 20.6 (q), 19.6 (t), 18.3 (q), 13.8 (q). MS (ESI^+^), *m/z* (*I_rel_*, %): 1307 [2M+Na]^+^ (12), 643 [M+H]^+^ (100).

*3β-Acetoxy-5α,8α-(1,4-dioxo-1,2,3,4-tetrahydrophthalazine-2,3-diyl)-27-norcholest-6-ene-25-one (***6***).* A mixture of pulverized NiCl_2_ × H_2_О (833 mg, 3.5 mmol), Zn powder (910 mg, 14 mmol), and but-3-en-2-one (285 μl, 3.5 mmol) in dry pyridine **(**20 ml) was stirred at 60 °C under argon for 30 min. The resulting dark red complex was cooled to room temperature and added to a solution of 22-iodides **5** (450 mg, 0.70 mmol**)** in dry pyridine (4 ml). The reaction mixture was stirred at room temperature for 2 hours, diluted with EtOAc, and then filtered through Celite. The organic phase was washed with 1 M HCl (60 ml) and brine (40 ml), dried over Na_2_SO_4_, the solvent was evaporated, and the residue was purified by column chromatography on silica gel (petroleum ether/EtOAc = 15:1) to give 25-ketone **6** (312 mg, 76%) as a light yellow oil. IR (film) ν, cm^−1^: 2955, 1755, 1735, 1715, 1310, 1250. ^1^H NMR (CDCl_3_, 500 MHz): δ_H_ 8.11 (m, 2H), 7.68 (m, 1H), 6.64 (d, *J* = 7.9 Hz, 1H), 6.28 (d, *J* = 7.9 Hz, 1H), 4.69 (m, 1H), 3.99 (dd, *J* = 14.1, 3.8 Hz, 1H), 3.91 (dd, *J* = 14.1, 3.8 Hz, 1H), 2.12 (s, 3H), 2.00 (s, 3H), 1.03 (s, 3H), 0.93 (d, *J* = 6.2 Hz, 3H), 0.81 (s, 3H). ^13^C NMR (CDCl_3_, 125 MHz): δ_C_ 209.2 (s), 170.1 (s), 162.1 (s), 159.7 (s), 137.7 (d), 132.8 (d), 132.8 (d), 130.5 (s), 129.9 (s), 129.0 (d), 127.0 (d), 126.9 (d), 69.9 (d), 68.2 (s), 66.8 (s), 56.4 (d), 50.3 (d), 48.7 (d), 44.3 (s), 44.2 (t), 40.4 (s), 39.2 (t), 35.1 (d), 35.0 (t), 34.9 (t), 31.1 (t), 29.9 (q), 27.6 (t), 26.0 (t), 24.4 (t), 21.7 (t), 21.3 (q), 20.3 (t), 18.4 (q), 18.3 (q), 13.1 (q). MS (ESI^+^), *m/z* (*I_rel_*, %): 587 [M+H]^+^ (100).

*3β-Acetoxy-5α,8α-(1,4-dioxo-1,2,3,4-tetrahydrophthalazine-2,3-diyl)-25-hydroxycholest-6-ene (***7***).* Methylmagnesium iodide in Et_2_O (1 M, 2 ml, 2 mmol) was added to a stirred and cooled to 0 °C solution of 25-ketone **6** (312 mg, 0.53 mmol) in dry Et_2_O (10 ml) for 5 min under argon. The reaction mixture was stirred at ice bath temperature for 15 min and at room temperature for 2 h. Then, the reaction mixture was carefully quenched with saturated NH_4_Cl (8 ml), diluted with EtOAc (10 ml), washed with brine (30 ml), and dried over Na_2_SO_4_. The solvent was evaporated, and the residue was purified by column chromatography on silica gel (petroleum ether/EtOAc = 12:1) to give 25-alcohol **7** (154 mg, 48%) as a light yellow oil. IR (film) ν, cm^−1^: 3450, 2970, 2855, 1755, 1735, 1310, 1250. ^1^H NMR (CDCl_3_, 500 MHz): δ_H_ 8.13 (m, 2H), 7.70 (m, 1H), 6.66 (d, *J* = 7.9 Hz, 1H), 6.29 (d, *J* = 7.9 Hz, 1H), 4.70 (m, 1H), 4.00 (dd, *J* = 14.1, 6.9 Hz, 1H), 3.92 (dd, *J* = 14.1, 6.9 Hz, 1H), 2.01 (s, 3H), 1.22 (s, 6H), 1.04 (s, 3H), 0.94 (d, *J* = 6.4 Hz, 3H), 0.83 (s, 3H). ^13^C NMR (CDCl_3_, 125 MHz): δ_C_ 170.2 (s), 162.2 (s), 159.7 (s), 137.7 (d), 132.9 (d), 132.8 (d), 130.5 (s), 129.9 (s), 129.1 (d), 127.1 (d), 126.9 (d), 70.1 (s), 69.9 (d), 68.3 (s), 66.9 (s), 56.4 (d), 50.3 (d), 48.7 (d), 44.3 (s), 44.2 (t), 40.4 (s), 39.4 (t), 35.1 (t), 35.0 (d), 34.9 (t), 31.1 (t), 30.6 (q), 30.4 (q), 27.6 (t), 26.1 (t), 24.5 (t), 21.7 (t), 21.4 (q), 20.4 (t), 18.5 (q), 18.3 (q), 13.1 (q). MS (ESI^+^), *m/z* (*I_rel_*, %): 603 [M+H]^+^ (36), 585 [M–Н_2_О+H]^+^ (100).

*25-hydroxy-7-dehydrocholesterol (25OH7DHC, (***8***).* To a solution of LiAlH_4_ (99 mg, 2.60 mmol) in dry THF (3 ml), a solution of 25-ketone **7** (154 mg, 0.26 mmol) in dry THF (3 ml) was added dropwise at 0 °C under argon. The reaction mixture was refluxed for 2 h, diluted with EtOAc, washed with 0.5 M HCl (10 ml), NaHCO_3_ (saturated, 10 ml), and brine (20 ml), and then dried over Na_2_SO_4_. The solvent was evaporated, and the residue was purified by column chromatography on silica gel (petroleum ether/EtOAc = 8:1) to give 25OH7DHC **8** (81 mg, 48%) as a white solid; mp 157-159 °С. UV (MeOH) l_max_, nm (e): 272 (8453), 282 (9136), 293 (6155). IR (KBr) ν, cm^−1^: 3390, 2970, 2940, 2860, 1070. ^1^H NMR (C_5_D_5_N, 500 MHz): δ_H_ 5.72 (d, *J* = 3.4 Hz, 1H), 5.52 (d, *J* = 2.1 Hz, 1H), 3.97 (m, 1H), 2.83 (d, *J* = 14.4 Hz, 1H), 2.70 (d, *J* = 14.4 Hz, 1H), 1.45 (s, 6H), 1.06 (s, 3H), 1.03 (d, *J* = 6.4 Hz, 3H), 0.67 (s, 3H). ^13^C NMR (C_5_D_5_N, 125 MHz): δ_C_ 141.7 (s), 141.4 (s), 120.1 (d), 117.5 (d), 70.3 (d), 70.0 (s), 56.6 (d), 55.1 (d), 47.0 (d), 45.7 (t), 43.6 (s), 42.4 (t), 39.9 (t), 39.4 (t), 37.9 (s), 37.5 (t), 36.9 (d), 33.4 (t), 30.6 (q), 30.4 (q), 28.8 (t), 23.8 (t), 21.8 (2t), 19.5 (q), 17.0 (q), 12.4 (q). MS (ESI^+^), *m/z* (*I_rel_*, %): 383 [M–H_2_O+H]^+^ (100), 365 [M–2H_2_O+H]^+^ (40).

**Synthesis of (E)-N**'**-hydroxy-N-(4-isopentyl-2-methylphenyl)formimidamide (cpd5')*.***

The scheme of synthesis is shown in Fig. S8. The synthesis was performed in 5 steps.

*1.	2-methyl-4-(3-methylbut-1-en-1-yl)-1-nitrobenzene (****2***'*):*

The ground isobutyltriphenylphosphonium bromide (281.2 g, 0.7 mol) and toluene (1600 ml) were placed under positive flow of argon in a dry 4-liter four-neck round-bottom flask equipped with an overhead mechanical stirrer, dropping funnel, and thermometer. Solution of potassium t-butoxide (72.2 g, 0.71 mol) in THF (380 ml) was added to the mixture via the dropping funnel for 5 min. The funnel was rinsed with 20 ml of THF after the addition, and the red reaction mixture was stirred for 3.5 h at room temperature. Then, the solution was cooled to –78 °C and aldehyde (**1)** (90 g, 0.55 mol) in a mixture of toluene (250 ml) and THF (50 ml) was added dropwise over 45 min, keeping the temperature below –70 °C. The funnel was rinsed with 20 ml of THF after the addition. The mixture was allowed to warm to room temperature, and it was stirred overnight. The reaction was quenched with aqueous NH_4_Cl (36 g in 500 ml water), and the obtained mixture was then stirred for 15 min. The organic layer was then separated, washed with H_2_O (300 ml), dried over Na_2_SO_4_, filtered, and concentrated in vacuo. The resulting residue was diluted with petroleum ether (1000 ml), filtered, and the precipitate (triphenylphosphine oxide) was additionally washed with hot petroleum ether (two times, 500 ml). The combined filtrates were concentrated under reduced pressure, and the resulting residue was dissolved in dichloromethane and treated with 9 ml of 30% hydrogen peroxide solution. The mixture was stirred overnight at room temperature, and then sodium sulphite (7.4 g) was added followed by sodium sulphate during 30 min of stirring. The precipitate was filtered off and washed with dichloromethane (100 ml). The combined filtrates were concentrated under reduced pressure, and the resulting residue was diluted with petroleum ether (250 ml). The formed precipitate was filtered off and washed with petroleum ether (50 ml). The combined filtrates were concentrated under reduced pressure, and the obtained residue was chromatographed on silica gel (petroleum ether: ethyl acetate, 100:0 to 98:2) to afford alkene (**2)** (105 g, 93%) as mixture of *cis*- and *trans*-isomers.

^1^H NMR (500 MHz, CDCl_3_, signals of main isomer) δ_H_ 7.98 (d, *J* = 8.4 Hz, 1H), 7.21 (d, *J* = 8.5 Hz, 1H), 7.18 (s, 1H), 6.27 (d, *J* = 11.7 Hz, 1H), 5.67 – 5.58 (m, 1H), 2.83 (qd, *J* = 13.1, 6.6 Hz, 1H), 2.62 (s, 3H), 1.06 (d, *J* = 6.6 Hz, 5H).

*2.	4-isopentyl-2-methylaniline (****3***'*):*

A suspension of Raney nickel (10 g, washed twice with ethanol) and compound **(2**'**)** (90 g, 0.439 mol) in ethanol (450 ml) was stirred at 50 ºC under a hydrogen atmosphere until both nitro group and alkenyl groups were completely reduced (36-48 h, control: NMR). The catalyst was separated by decanting and washed with ethanol (3 times, 50 ml). The combined filtrates were concentrated, and the residue was filtered through a short plug with sodium sulphate (with toluene as the eluent) to afford compound (3') (70 g, 90%) after concentration under reduced pressure.

^1^H NMR (500 MHz, CDCl_3_) δ_H_ 6.94 (s, 1H), 6.91 (d, *J* = 8.1 Hz, 1H), 6.68 (d, *J* = 7.9 Hz, 1H), 3.77 (br.s, 2H), 2.55 (dd, *J* = 14.8, 6.8 Hz, 2H), 2.22 (s, 3H), 1.68 – 1.57 (m, 1H), 1.54 – 1.46 (m, 2H), 0.97 (d, *J* = 6.6 Hz, 6H).

*3.	N*'*-(4-isopentyl-2-methylphenyl)-N,N-dimethylformimidamide (****4***'*):*

A solution of compound **3**' (70 g, 0.395 mol) and N,N-dimethylformamide dimethyl acetal (75 ml, 0.562 mol) in dry toluene (300 ml) was refluxed for 4 hours under positive nitrogen pressure. The mixture was cooled and concentrated under reduced pressure. The resulting residue (91 g) was used for the next step without purification.

*4.	N*'*-hydroxy-N-(4-isopentyl-2-methylphenyl)formimidamide (****5***'*):*

NH_2_OH-HCl (33 g, 0.479 mol) was added to a solution of crude **4**' (70 g, 0.3 mol) in ethanol (1500 ml), and the mixture was stirred at room temperature for 2 hours and concentrated under reduced pressure. Water (100 ml) and dichloromethane (600 ml) were added to the residue and the resulting mixture was vigorously stirred for 5 min, after which the organic layer was separated, washed with brine, and dried over sodium sulphate. The dried solution was concentrated, and the resulting residue was crystallized from a mixture of petroleum ether (450 ml) and iso-propanol (50 ml) at –18 ºC during 8 hours. The precipitate was filtered off, washed with petroleum ether, and dried at room temperature. The mother liquor and washings were combined, evaporated, and the resulting residue was dissolved in a mixture of petroleum ether (45 ml) and iso-propanol (5 ml) to perform a second crystallization as described above. The total yield of the crystallized product (purity >98%) was 57.4 g (85% based on aniline (**3**')). mp 125-126 ºС.

^1^H NMR (500 MHz, CDCl_3_) δ_H_ 7.36 (s, 1H), 7.08 – 6.96 (m, 3H), 6.93 (d, *J* = 8.7 Hz, 1H), 6.43 (s, 1H), 2.62 – 2.45 (m, 2H), 2.26 (s, 3H), 1.64 – 1.52 (m, 1H), 1.47 (dd, *J* = 15.6, 7.2 Hz, 2H), 0.93 (d, *J* = 6.6 Hz, 6H).

^13^C NMR (125 MHz, CDCl_3_) δ_H_ 140.90 (d), 137.83 (s), 135.43 (s), 131.24 (d), 127.13 (d), 125.81 (s), 115.23 (d), 41.13 (t), 33.11 (t), 27.80 (d), 22.67 (q), 17.47 (q).
